# Optimal immune specificity at the intersection of host life history and parasite epidemiology

**DOI:** 10.1101/2021.03.11.434955

**Authors:** A. E. Downie, A. Mayer, C. J. E. Metcalf, A. L. Graham

## Abstract

Epidemiology and life history are commonly hypothesized to influence host immune strategy, and the pairwise relationships between immune strategy and each factor have been extensively investigated. But the interaction of these two is rarely considered, despite evidence that this interaction might produce emergent effects on optimal immune strategy. Here we investigate the confluence of epidemiology and life history as it affects immune strategy through a demographically-framed model of sensitivity and specificity in parasite recognition and response. We find that variation in several different life history traits associated with both reproduction and longevity alters optimal immune strategies – but the direction and magnitude of these effects depends on how epidemiological risks vary across life. Drawing on published life history data, we also find that our predictions apply across chordate taxa. Our results shed light on the complex interactions shaping immune strategy and may prove valuable in interpreting empirical results in ecoimmunology.

## Introduction

Parasites are a central threat to organismal fitness, responsible for a vast share of mortality. Accordingly, all organisms possess some anti-parasite capacities, which can be brought together under the umbrella of immune defenses. We expect immune defenses to be tailored to a host’s ecological context to maximize efficacy and efficiency, particularly since immune defense incurs resource, immunopathology, and other costs (Sheldon and Verhulst 1996, Lochmiller and Deerenberg 2000, Zuk and Stoehr 2002, Schmid-Hempel 2003). Accordingly, the strategies hosts use may differ. For example, should parasites be directly resisted to extirpate them, or should their damage be tolerated and contained without directly countering the parasites (Råberg et al. 2008, Medzhitov et al. 2012)? Should immune investment be constitutive or facultative (Schmid-Hempel 2003)? Should parasite recognition systems be very sensitive to possible threats or highly conservative and specific in identifying them (Metcalf et al. 2017)? Different strategies imply deployment of different immune defenses and in different quantities. In principle this applies to both inter- and intraspecific comparisons – there should be variation in immune strategies within populations and across species. These strategic conundrums accordingly offer a promising path for developing predictions for how organisms might differ in their immune defenses and disease ecology.

But what factors might shape immune strategy? One commonly-hypothesized ecological factor affecting immune strategy and deployment is epidemiology (Shudo and Iwasa 2001, Hamilton et al. 2008, Lazarro and Little 2009, Boots et al. 2009, Donnelly et al. 2015). For example, we expect investment in immune defenses to rise with increasing parasite threat (Hamilton et al. 2008) – without parasites, why waste resources on immune defense? Furthermore, we expect variability and diversity in parasite threats to affect the apportionment of that investment to different defenses (Shudo and Iwasa 2001, Hamilton et al. 2008, Mayer et al. 2016) and the architecture of those defenses (Kubinak et al. 2012, Mayer et al. 2015, Chen et al. 2021). Immunopathology – damage inflicted on the self by the immune system – also presents a significant risk to hosts; balancing immunopathology with averting parasite-inflicted damage may constrain investment in immune defenses and alter their form (Graham et al. 2005, Cressler et al. 2015, Metcalf et al. 2017). While assessing details of epidemiological risk in empirical settings can be difficult, empirical work has generally supported the influence of disease environment on optimal immune defense and strategy (Duffy et al. 2012, Westra et al. 2015, Chen et al. 2021).

Host life history has also been proposed to affect immune strategy (Sheldon and Verhulst 1996, Zuk and Stoehr 2002, Ricklefs and Wikelski 2002, Lee 2006, Valenzuela-Sánchez et al. 2021). This idea springs from the aforementioned resource costs of immunity, since life history is intimately tied to resource allocation (Stearns 1992). Correspondingly, immune defenses have been proposed to be linked to the fast/slow spectrum of life history and the related pace-of-life concept (Ricklefs and Wikelski 2002, Lee 2006). In particular, immune investment is predicted to correlate negatively with reproductive output (Sheldon and Verhulst 1996, Zuk and Stoehr 2002, Lee 2006) and positively with longevity (Lee 2006). Furthermore, Lee (2006) suggests that species with a faster life history may use more specific immune defenses and fewer inflammatory immune defenses. Some experimental evidence supports a negative trade-off between immune investment and reproduction (Knowles et al. 2009), but generally empirical investigations of the relationship between immune strategy and life history have been inconclusive or inconsistent (e.g. Martin et al. 2007, Lee et al. 2008).

While epidemiology and life history are both considered important for immune defense, their intersection has rarely been explored. The few studies that exist have uncovered intriguing contingencies for the relationship between immunity, epidemiology, and life history. For example, Donnelly et al. (2017) have shown that the diversity of parasites in a host’s environment can shape the relationship of lifespan with immune memory. In one-parasite systems, optimal investment in memory peaks at intermediate lifespans due to ecological feedbacks; essentially, herd immunity, by suppressing parasite transmission, creates opportunities for low-memory strategies to succeed (Van Boven and Weissing 2004, Miller et al. 2007, Boots et al. 2009, Best and Hoyle 2013, Boots et al. 2013). In two-parasite systems, that effect disappears, and optimal memory investment increases with increasing lifespans (Donnelly et al. 2017). Thus the epidemiological context (here the diversity of parasites) shapes the relationship between life history (lifespan) and optimal immune strategy (immune memory).

These results suggest that life history and epidemiology interact to determine optimal immune strategies and may thereby explain why strategies vary. Here we are especially interested in variation in epidemiological environment with age. Empirical work shows that infection risk and burden vary across life, with different profiles of age-dependence for different host species (Hudson and Dobson 1995, Wilson et al. 2002). For example, in Cape buffalo (*Syncerus caffer*), only some of its parasites infect it during the first year of life (Combrink et al. 2020). And sexually transmitted infections will generally only pose a threat to hosts after reproductive maturity (Antonovics et al. 2011). Immune strategies for some taxa may be pulled in multiple directions by different infection threats at different stages of life. Therefore, by studying the intersection of age-dependent epidemiological variation with life history variation, we aim to elucidate causes of varied immune strategies across host taxa.

To this end we extend the model of immune sensitivity and specificity developed in Metcalf et al. (2017) and Metcalf and Graham (2018). The logic of the basic model is as follows. Hosts face a variety of antigenic stimuli, from harmful sources – generally non-self – and benign sources. The host must correctly ascertain from these stimuli whether or not it is infected and should mount an immune response (Fig. 1). As in other signal detection systems (Nesse 2005), a trade-off between sensitivity and specificity then arises: an organism with a more sensitive immune system will more frequently correctly identify parasite stimuli and attack them, but it will also more often produce immunopathology by mistaking self molecules for parasite signals. An organism that is more specific is less likely to produce a damaging autoimmune response, but it is also more likely to ignore a parasite and therefore suffer more parasite-inflicted damage. Sensitivity and specificity might find expression in receptor structure or in activation thresholds for the immune system (Metcalf et al. 2017). Combining sensitivity and specificity with epidemiological risks – infection risk, infection mortality risk, and immunopathology risk – allows us to determine survival in a given risk environment (Fig. 1A). We can then use a demographic matrix framework to provide resolution on both life history and epidemiological variation. The optimal immune strategy is that which maximizes fitness; we define fitness here as the population growth rate, λ, which describes the rate of increase through time. Prior work has shown that the relative balance of these different types of epidemiological threats shapes immune strategy: in general, an organism facing greater infection risk should be more sensitive, and an organism facing lower infection risk should be more specific (Metcalf et al. 2017, Metcalf and Graham 2018). Background mortality may also influence optimal defense strategy, but only when disease or immunopathology risks vary with age (Metcalf et al. 2017, Metcalf and Graham 2018). When such risks do not vary with age, then no aspect of demography affects immune strategy (Metcalf and Graham 2018). However, as discussed above, such risks are rarely static.

**Figure 1:**
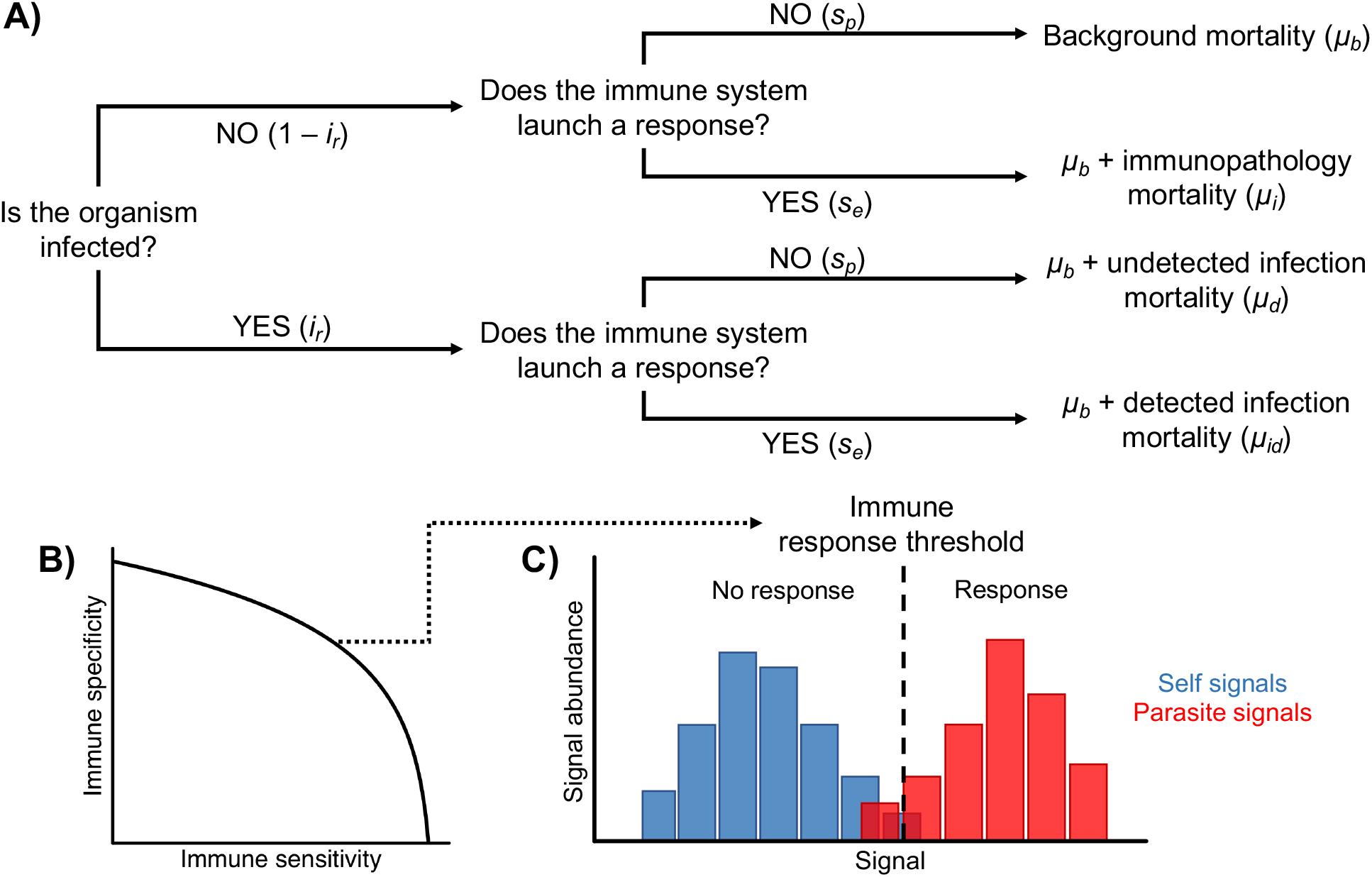
Sensitivity and specificity in immunity. A) Decision tree outlining the relationship of sensitivity and specificity to infection and mortality risks. Sensitivity and specificity, together with frequency of infection, determine the relative balances of various types of mortality. B) Schematic showing how sensitivity and specificity manifest in response to distributions of benign (blue) and pathogenic (orange) signals. Different combinations of sensitivity and specificity define the location of the response threshold. A more sensitive immune system has a response threshold shifted to the left, responding to a greater range of signals; a more specific immune system has a response threshold shifted to the right, responding to a smaller range of signals. C) Receiver-operator curve showing trade-off between sensitivity and specificity in immunity. Different points on trade-off curve indicate different combinations of sensitivity and specificity and produce different response thresholds. The shape of the trade-off curve is governed by how much host and parasite signals overlap and the capacity of the host to discern fine molecular differences.

Here we explicitly study the interaction between life history and epidemiology as it influences optimal immune strategy, drawing on a variety of scenarios reflecting empirical patterns. We mix variation in life history strategies with variation in epidemiological risk across life. When we refer to epidemiology in our approach, we are referring to the risk of being infected and the mortality risks associated, rather than the details of disease dynamics. First we focus on different simple reproductive output schedules, exploring how they shape optimal immune specificity and sensitivity in various epidemiological contexts. We then marry a range of realized life histories drawn from the COMADRE database of demographic matrices (Salguero-Gómez et al. 2016) with epidemiological risk schedules to explore in greater detail the complex interplay between life history, epidemiology, and immune strategy. We find that this confluence is indeed critical, with epidemiology mediating both the strength and direction of the relationship between life history traits and optimal immune strategy.

## Results

### Reproductive Demography Affects Optimal Immune Strategy

We explored five different schedules for reproduction in our analysis of the relationship between reproductive demography and optimal immune strategy (Fig. 2). For a given set of epidemiological risk parameters, each reproductive schedule has a different associated optimal immune sensitivity and specificity maximizing λ (Fig. 2). In the case considered here, where infection risk drops at the age of first reproduction, a life history associated with higher reproductive output leads to lower sensitivity and higher specificity, and vice-versa for lower reproductive output. The two reproductive schedules having changes in output with age are further polarized in their associated optimal immune strategies (Fig. 2). Each reproductive schedule also associates with different stable age structures and distributions of reproductive value and elasticity of λ with respect to survival and fecundity (Fig. S3). Because different reproductive schedules produce different values of λ, we also explored optimal immune strategies when we manipulate *μ_b_* to equalize λ across the reproductive schedules; the effect of reproductive schedule on immune strategy is unaltered (Table S3). Thus it is not λ that is associated with a given immune strategy, it is the reproductive schedule itself.

**Figure 2:**
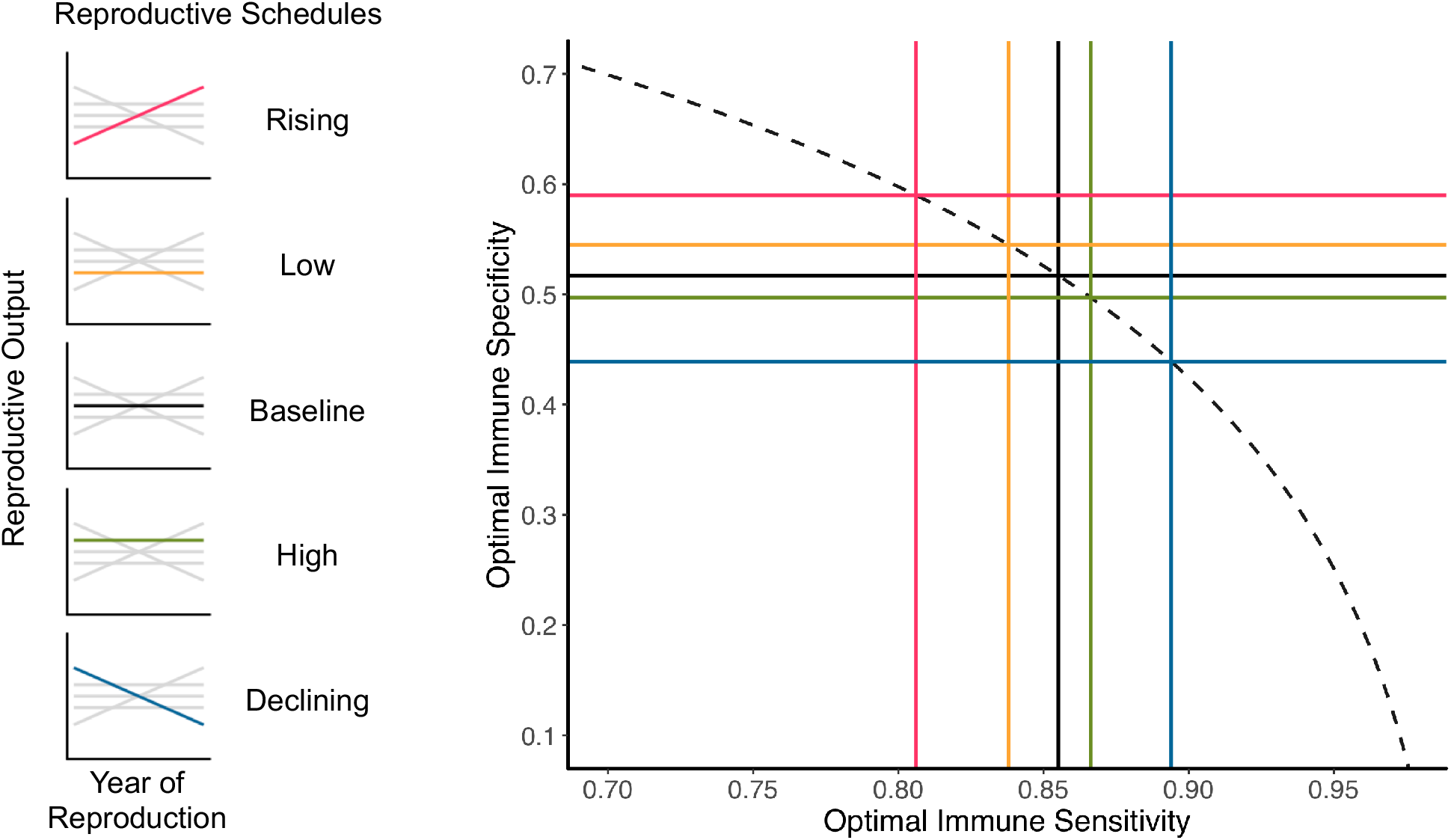
Influence of reproductive demography on optimal immune strategy. Plot showing optimal combination of immune sensitivity and specificity for each of five reproductive demographic schedules (at left), for an epidemiological risk environment where infection risk *i_r_* = 0.6 for the first two age classes (prior to reproductive maturity) and *i_r_* = 0.2 thereafter. Reproduction begins in the third age class for all demographic schedules. Strategy optima are determined as the immune specificity and sensitivity maximizing λ, the population growth rate. Dashed curve shows the shape of the specificity/sensitivity trade-off curve for γ = 4. Other parameter values are *μ_b_* = 0.15, *μ_i_* = 0.1, *μ_d_* = 0.3, and *μ_id_* = 0.01.

### Epidemiological Risk Environment Modulates Effects of Reproduction on Immune Strategy

Different organisms experience different epidemiological risks across their lives. We therefore expanded on the analysis above by exploring how the relationship between reproductive demography and immune strategy changes in different epidemiological scenarios (Fig. 3). We find that a different epidemiological context can completely reverse the direction of this relationship (Fig. 3A). In an epidemiological setting with *i_r_* decreasing at reproductive maturity, the late-skewed rising reproductive schedule (red) has the most specific immune strategy of our five reproductive schedules, whereas when *i_r_* rises at reproductive maturity, that same reproductive schedule associates with the least specific immune strategy. The early-skewed declining reproductive schedule (blue) has the opposite set of associations in these two epidemiological environments. In essence, a “flip” in the ordinal relationship between reproductive and immune strategies takes place when the epidemiological schedule is reversed. This reversing effect also appears when we explore variation in *μ_d_* across life while holding *i_r_* constant (Fig. S4A).

**Figure 3:**
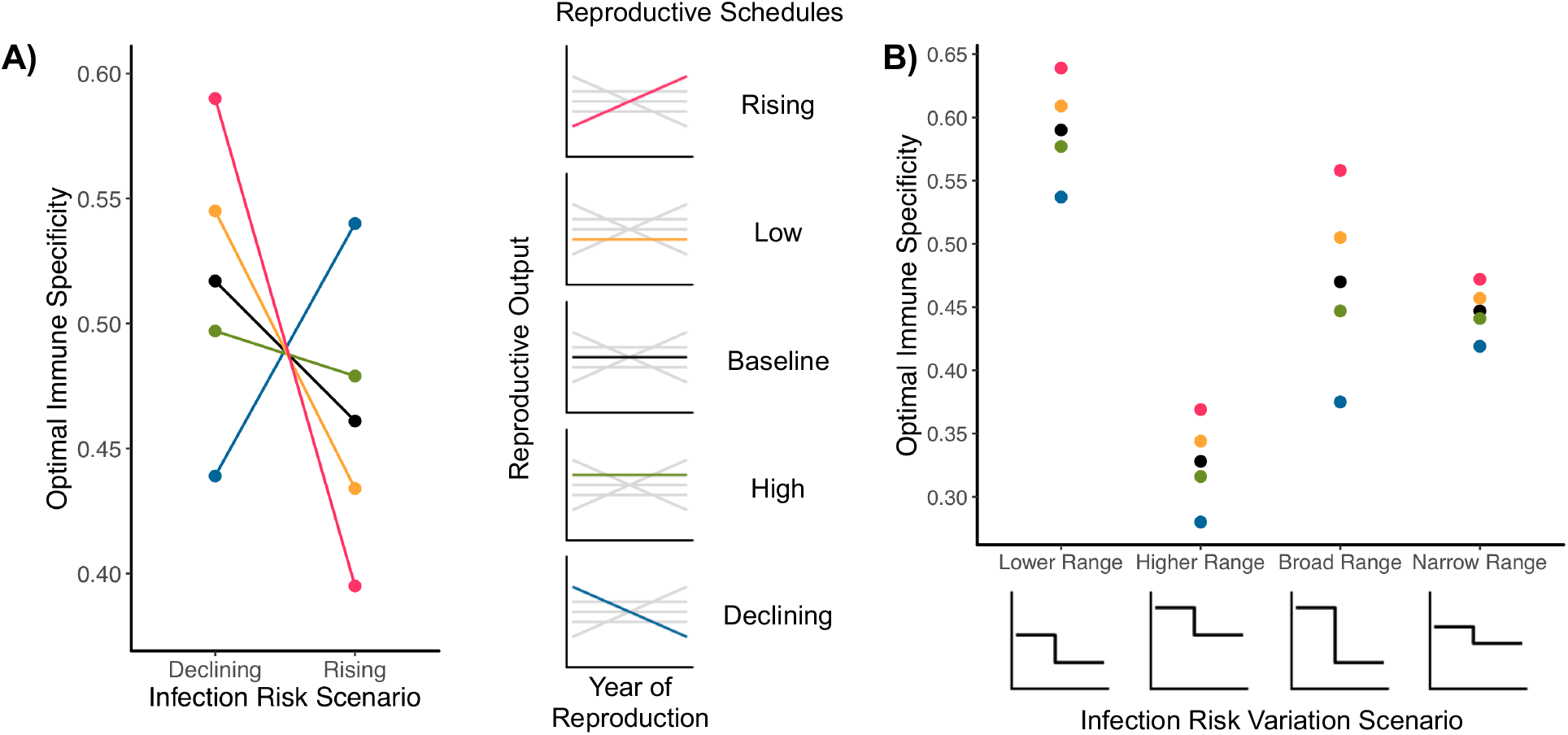
Epidemiological environment alters the effect of reproduction on immune strategy: infection risk *i_r_*. Reproduction begins in the third age class for all schedules. In each scenario, *i_r_* starts at one value and changes to a different value at the third age class. A) The change in optimal immune specificity associated with differences in epidemiological context (i.e. changes in *i_r_*, on the x-axis) and reproduction (different points and lines). In the declining scenario, *i_r_* declines drops at reproductive maturity from 0.6 to 0.2; in the rising scenario, *i_r_* increases from 0.2 to 0.6. Other parameter values are *μ_b_* = 0.15, *μ_i_* = 0.1, *μ_d_* = 0.3, *μ_id_* = 0.01, and γ = 4. B) The change in range of optimal specificities associated with different reproductive demographies for different magnitudes of variation in decline of infection risk *i_r_* at reproductive maturity. In the lower range scenario, *i_r_* drops from 0.45 to 0.2; in the higher range, from 0.7 to 0.45; in the broad range, from 0.7 to 0.2; in the narrow range, from 0.525 to 0.375. Other parameter values are *μ_b_* = 0.15, *μ_i_* = 0.1, *μ_d_* = 0.3, *μ_id_* = 0.01, and γ = 4.

Differences in the range and magnitude of epidemiological risk variation with age alter the differences in optimal immune strategy between reproductive demographies. Overall level of risk produces different immune strategies, and when the magnitude of variation across life changes, the strength of the reproduction-immunity relationship changes (Fig. 3B). Here, greater *i_r_* variation across life is associated with greater differences in immune strategy between reproductive schedules. Variation in other risk parameters produces the same insight: when we examined variation in *μ_d_*, we found the same correlation between the strength of the reproduction-immune strategy relationship and degree of epidemiological risk variation (Fig. S4B). Overall, our results suggest we cannot know from life history alone which of two populations or species should be more sensitive or specific in immune defense without knowing the epidemiological context of each population. The epidemiology is essential.

### Epidemiology Interacts with Several Demographic Traits to Influence Immune Strategy

To expand on our above results, we considered the relationship between specific life history traits and optimal immune strategy. We did this by exploiting the life histories recorded in the COMADRE database of animal demographic matrices (Salguero-Gómez et al. 2016, COMADRE Animal Matrix Database 2020). These matrices provide reproductive schedules and estimated mortality curves across a wide taxonomic range, and we processed the raw matrices to mesh them with our immune strategy estimation approach without changing the essential demographic patterns encoded in said matrices (see Methods, Fig. S1). We combined the output matrices with various epidemiological scenarios to explore the change in predicted optimal immune strategy among different demographies in a given epidemiological context. We analyzed the relationship between predicted optimal immune specificity and life history traits with a Bayesian linear model. For epidemiological variation we used stepped variation in *i_r_* as above, where *i_r_* changes once at reproductive maturity.

We find correlations between predicted optimal immune strategy and three different life history traits – age class of first reproduction (AFR; approximating age of first reproduction), mean reproductive rate (MRR), and reproductive life expectancy (RLE) (Figs. 4, 5, Table 1). And we identify the same epidemiology-induced flip in the life history-immune strategy relationship that we describe above. The effects of the life history traits are interlocking, such that variation unexplained by one life history trait may be explained by another. An early age of first reproduction can be associated with a wide range of optimal immune strategies, but these strategies are in turn shaped by the mean reproductive rate (Fig. 4). For decreasing infection risk with age and an early age of first reproduction, high specificity corresponds to low rates of reproduction, and vice-versa. At later ages of first reproduction, the range of associated reproductive rates contracts, and optimal specificity becomes less variable and confined to relatively higher values (Fig. 4A). The reverse pattern is observed when infection increases with age (Fig. 4B, Table 1).

**Figure 4:**
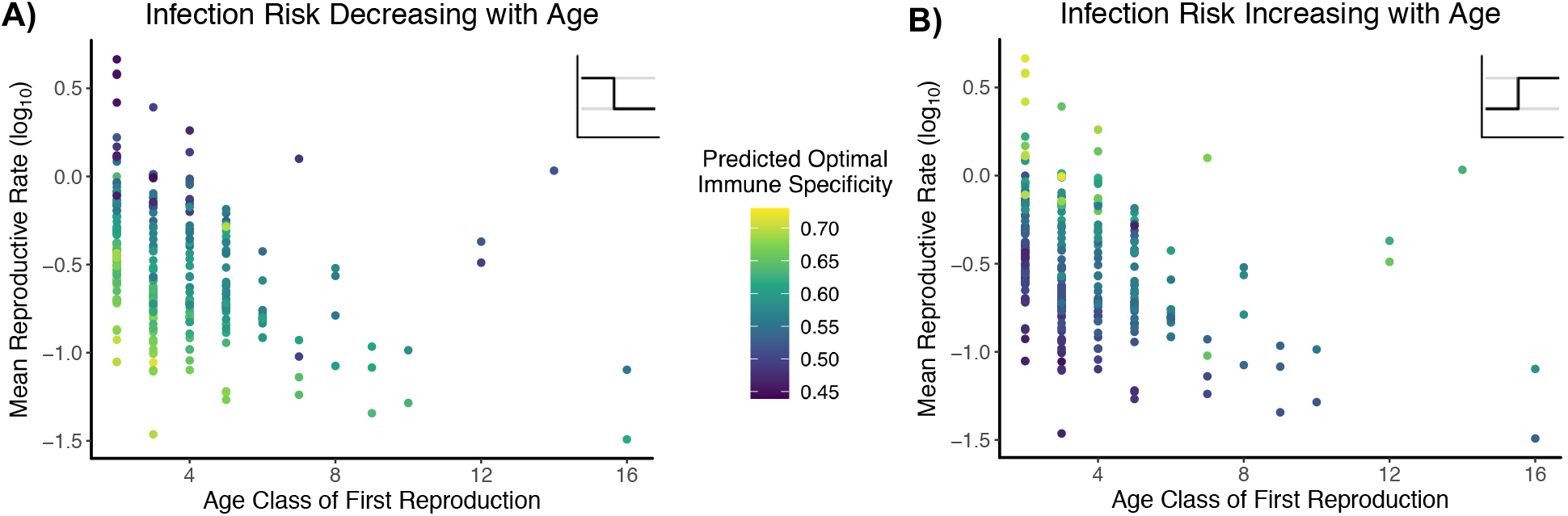
Interaction of demography and epidemiology for immune specificity. Infection risk is set such that there is one infection risk *i_r_* for age classes prior to reproductive maturity and a different risk *i_r_* for reproductive age classes. Our dataset comprises 298 population matrices representing 129 chordate species. For all scenarios, parameter values are *μ_d_* = 0.3, *μ_i_* = 0.1, *μ_id_* = 0.01, ρ = 0.75, and γ = 4. A) Predicted optimal immune specificities when infection risk declines with age, with respect to population age class of first reproduction and mean reproductive rate as calculated from original matrix. Infection risk *i_r_* prior to reproductive maturity is 0.45; for reproductive age classes, it is 0.2. B) Predicted optimal immune specificities when infection risk rises with age with respect to population age class of first reproduction and mean reproductive rate as calculated from original matrix. Infection risk *i_r_* prior to reproductive maturity is 0.2; for reproductive age classes, it is 0.45.

**Figure 5:**
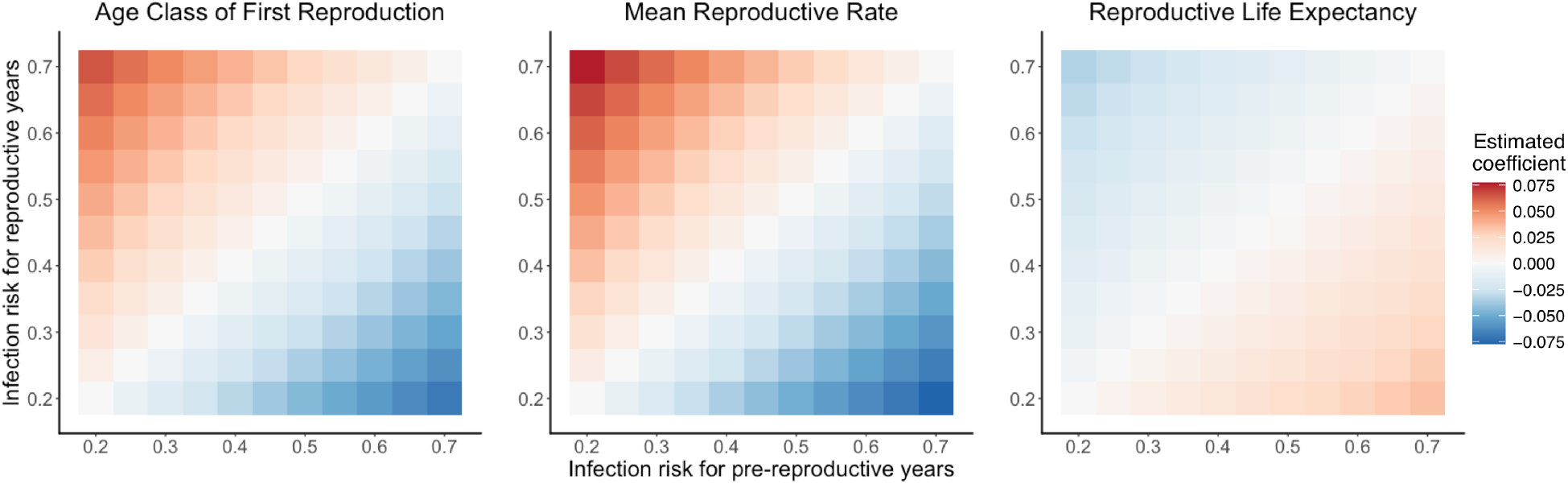
Epidemiological dependency of life history-immune specificity relationship. Tile plots showing, for a variety of scenarios of variation in *i_r_*, coefficients of relationship between predicted optimal immune specificity and designated life history trait as estimated from Bayesian linear models. Each tile is a scenario in which predicted optimal immune specificities were generated for 298 population projection matrices representing 129 chordate species. A linear model was used to estimate relationship coefficients for each scenario. Coefficient value in plot represents mean value of the posterior probability distribution, except on the diagonal; on the diagonal, all coefficients are 0 (see text). Value of *i_r_* on x-axis is the risk for pre-reproductive years, jumping in age class of first reproduction to value of *i_r_* on y-axis. Life history trait values were log-transformed and standardized as Z-scores for comparability of coefficients. For all estimated coefficients off the diagonal, 89% confidence intervals do not include 0. For all scenarios, parameter values are *μ_d_* = 0.3, *μ_i_* = 0.1, *μ_id_* = 0.01, ρ = 0.75, and γ = 4.

**Table 1:** Results from Bayesian linear model for demography and immune specificity. Linear model looks at predicted optimal immune specificity as a function of three different life history summary statistics. Results are means and, in brackets, boundaries of 89% highest posterior density intervals (HPDI) for posterior probability distributions. Entries in *italics* indicate the 89% HPDI overlaps with 0. All variables calculated from original matrix in COMADRE database, log-transformed, and standardized as Z-scores. Dataset includes 298 qualifying matrices from 129 chordate species. For stepped epidemiological scenario, when infection risk is rising, *i_r_* in pre-reproductive years is 0.2, and *i_r_* in reproductive years is 0.45. When infection risk declines in the stepped scenario, *i_r_* in pre-reproductive years is 0.45, and *i_r_* in reproductive years is 0.2. In smoothed declining scenario, *i_r_* declines from 0.45 to 0.2; in rising scenario, *i_r_* rises from 0.2 to 0.45. Other parameter values are *μ_d_* = 0.3, *μ_i_* = 0.1, *μ_id_* = 0.01, *ρ* = 0.75, and γ = 4.

| Parameter | Declining stepped infection risk $i_r$ | Rising stepped infection risk $i_r$ | Declining smoothed infection risk $i_r$ | Rising smoothed infection risk $i_r$ |
| --- | --- | --- | --- | --- |
| Intercept | 0.603<br>[0.600, 0.605] | 0.544<br>[0.541, 0.547] | 0.538<br>[0.536, 0.541] | 0.608<br>[0.605, 0.611] |
| Age class of first reproduction | -0.0381<br>[-0.0411, -0.0352] | 0.0357<br>[0.0327, 0.0384] | *$4.82 \times 10^{-4}$*<br>[*-0.00285, 0.00380*] | *-0.00121*<br>[*-0.00463, 0.00239*] |
| Mean reproductive rate | -0.0423<br>[-0.0454, -0.0390] | 0.0412<br>[0.0379, 0.0444] | -0.0409<br>[-0.0444, -0.0374] | 0.0440<br>[0.0402, 0.0479] |
| Reproductive life expectancy | 0.0190<br>[0.0155, 0.0224] | -0.0184<br>[-0.0218, -0.0150] | -0.0232<br>[-0.0269, -0.0197] | 0.0252<br>[0.0211, 0.0292] |
| Standard deviation | 0.0278<br>[0.0260, 0.0298] | 0.0280<br>[0.0260, 0.0300] | 0.0304<br>[0.0285, 0.0324] | 0.0322<br>[0.0301, 0.0344] |

In general, the strength and direction of the relationship between any given life history parameter and predicted optimal immune strategy depends on the level of infection risk and the amount it varies with age (Fig. 5). When risk does not vary at all with age, then there is no relationship between any individual demographic trait and immune specificity, regardless of absolute risk level (as previously described). However, as the magnitude of change in infection risk at reproductive maturity grows, the strength of each immunity-demographic trait relationship increases, with the sign of the relationship depending on which trait is being examined and whether risk increases or decreases at reproductive maturity. The estimated relationship between MRR and immune specificity is the strongest in all infection risk scenarios, followed by AFR and then RLE (Fig. 5, Table 1).

We further examined the robustness of our central finding with different datasets and other life history traits. The epidemiological dependency of the life history-immune strategy relationship is recapitulated when we repeat our analysis with a dataset of only mammal life histories (Table S4) and when we consider generation time as the sole life history trait influencing immune strategy in our linear model (Table S5). When we calculate our life history statistics from our matrices after dimension manipulation and mortality curve approximation (see Methods), results are qualitatively the same while differing to a small degree quantitatively (Table S6). This highlights that our matrix processing method, while producing matrices that approximate the original COMADRE matrices but are not identical, is not inducing significant distortions and generating mirages. All these analyses reinforce our main result, that epidemiological context determines the strength and direction of the relationship between life history traits and optimal immune strategy.

### Epidemiological Dependency of Life History-Specificity Relationship Spans Multiple Modes of Risk Variation

We also explored an alternate mode of infection risk variation with age. In this mode, rather than infection risk changing just once at reproductive maturity, we defined a risk for the first age class of the matrix and a risk for the last age class and allowed risk to change at a constant rate across intermediate age classes (see Methods, Fig. S2). This scenario represents a smooth change of infection risk with age. For use with our COMADRE process, we adjust the slope such that the relative change in infection risk with respect to matrix dimension (itself determined as a function of age at first reproduction and reproductive life expectancy) stays constant, ensuring comparability between matrices with different life histories (Fig. S2).

We repeated all of the preceding analyses for this second mode of infection risk variation, and we identified the same major results. Reproductive demography does influence optimal immune specificity and sensitivity (Fig. S5, Table S7), with the strength and direction of that influence depending on the direction and magnitude of change in epidemiological risk parameters (Figs. S6, S7). Furthermore, when we extend to our broader demographic analysis, we find the same epidemiological dependency for the strength and direction of the relationship between immune specificity and specific life history traits (Figs. S8, S9). Intriguingly, we do find different relative and absolute magnitudes for each relationship when compared to the stepped mode of infection risk variation (Tables 1, S4–S7, Fig. S9). If infection risk varies in this manner, our linear models are not confident that AFR has an influence on optimal immune strategy (Tables 1, S4–S7). And the direction of the relationships between optimal immune specificity and RLE are reversed (Fig. S9). This further highlights our finding that how epidemiological context varies with age shapes the relationship between life history traits and immune strategy. Particular demographic traits may have more or less influence on immune strategy, and in different directions, depending on precisely the epidemiological context.

## Discussion

Here, using a demographic model, we demonstrate that the influence of life history on immune strategy depends on the epidemiological context and the way that epidemiological risk varies across life. Both the strength and direction of the relationship, for a variety of different life history traits, are mutable. Thus, for example, when infection risk declines with age, a reproductive schedule with high output will push populations towards more sensitive and less specific immune strategies, while when infection risk rises with age, high reproductive output will select for less sensitive, more specific immune strategies. Infection risk variation could be created by factors like sociality (Altizer et al. 2003). In non-social species we might expect infection risk to be relatively low pre-reproduction, with a rise at reproductive maturity associated with the increased contact of mating and sexually-transmitted infections. In such species, those with greater reproductive output should have less sensitive, more specific immune strategies than those with lower reproductive output. But closely-related social species might not have the same variation in infection risk across age; perhaps they have a high frequency of early-life infections relative to infection in adulthood (see Altizer et al. (2003) for more discussion of sociality and infection). In such species, we would find that high reproductive output selects on immune strategy in the opposite direction, pushing for more sensitive, less specific defenses. We find a similar context-dependency for the relationship of immune strategy to age at first reproduction, a development- and longevity-associated trait, and reproductive lifespan, a longevity-associated trait. These results represent a novel insight into the importance of ecology for optimal immune strategy.

Our use of age classes allows us to effectively describe lifelong nuances in both life history and epidemiology, both of which vary among populations and species. Most prior studies of immune defense have used a constant background mortality to define lifespan. Such studies have produced intriguing results on their own (e.g. Best and Hoyle 2013, Donnelly et al. 2017, Metcalf et al. 2017), but two species with the same average or maximum lifespan may have radically different survival curves and therefore demographic patterns (Ronget and Gaillard 2020). By here describing lifespan in terms of age of first reproduction and reproductive life expectancy, we have identified how different facets of lifespan may actually select for different strategies of immune defense. For example, two species might have similar average lifespans but different ages at first reproduction and reproductive life expectancies. Our model tells us that we should expect them to differ in their immune strategy. This relationship between immune strategy and demography may also prove illuminating for the study of sex differences in immune defense (Zuk and Stoehr 2002, Metcalf and Graham 2018); males and females within a species may differ demographically – and in infection risks – in such a way as to push them towards different immune strategies. In general, our results show that greater nuance in life history and epidemiology offers greater insight into optimal immune defense.

One particularly intriguing result is that differences in reproductive schedule alone can influence optimal immune strategy when epidemiological risk varies with age. This surprising finding may be explainable by the relationship between the reproductive schedule and stable age structure and reproductive value in a population. Donnelly et al. (2015) show that optimal immune resistance should be tuned to favor stages that contribute the most to the rate of population growth (in our case λ). In our model, greater reproductive output leads to the elasticity of λ with respect to survival and fertility being relatively greater in early age classes and relatively lower in late age classes (Fig. S3C, D), indicating a relatively greater contribution to fitness in early age classes (Caswell 2001); accordingly, from Donnelly et al. (2015) we would expect an immune strategy that boosts survival in those early stages. And, indeed, that is what we find: when infection risk declines with age, greater reproduction associates with greater sensitivity to mitigate greater infection risk in early life. The rising and declining schedules produce the most divergent elasticity distributions, weighted towards late and early stages respectively, and accordingly have the most divergent immune strategies. And because we measure fitness through λ, the temporal aspect of population growth is critical. While the same immune strategy could maximize net reproductive rate R0 for the baseline, high, and low reproductive schedules, there are different immune strategies for each reproductive schedule to produce the fastest rate of population growth that ensures competitive success. Thus reproduction may influence optimal immune strategies through its effects on demographic structure in a population, not solely via the medium of resource allocation trade-offs (Knowles et al. 2009, Cressler et al. 2015).

Our results also suggest that theories predicting immune strategy primarily or only from pace-of-life or the fast-slow spectrum of life history (Ricklefs and Wikelski 2002, Lee 2006) may not be accurate, at least for immune sensitivity and specificity. There are two reasons for this. First, the direction of the association between any given life history trait and optimal specificity is contingent upon epidemiology, and thus reproduction or longevity may not necessarily always have the same association with immune strategy. Second, in certain epidemiological contexts, increased reproduction and increased longevity (which occupy opposite ends of the fast/slow spectrum) push immune specificity in the same direction. And AFR and RLE have opposite associations with immune specificity despite both being associated with a slow pace of life (Healy et al. 2019). Further work is necessary to establish whether this conclusion applies to other aspects of immune strategy, like resistance and tolerance. In particular, the aforementioned resource allocation trade-offs may influence different aspects of the immune system to greater or lesser degrees. Furthermore, particular life histories and ecological strategies may be associated with particular epidemiological risks (Albery and Becker 2020, Silk and Hodgson 2021), which may more closely tie life history to immune strategy. These are promising directions for future research.

Our findings may help explain why empirical investigations have struggled to consistently identify relationships between life history and immune defense when looking across multiple taxa. For example, lifespan correlates positively with immune investment in some (Tella et al. 2002) but not all studies (Nunn 2002, Martin et al. 2007, Downs et al. 2020) looking at various different suites of taxa. Our results here highlight two potential explanations for the rejection of this hypothesis in these studies. First, as described above, different aspects of lifespan produce different selective pressures on immune defense; thus the precise life histories of the species being compared are important context. As mentioned above, for cases where two organisms have similar overall projected lifespans but different ages at first reproduction and reproductive life expectancies, we would expect different their immune specificities. Second, in general these studies do not incorporate variation in epidemiological risk across taxa. The disease ecology of each species shapes strategy to a great extent, especially via level of risk (Hamilton et al. 2008, Donnelly et al. 2015, Westra et al. 2015, Metcalf et al. 2017). As long as epidemiological context varies – for example, infection risk rising with age in some species and declining with age in others – immune strategy would not correlate with life history across all of those species. None of this is to say that life history does not matter – we clearly find that it can matter a great deal. But researchers studying variation in immune strategy should consider and examine both epidemiology and demography, including variation in risk with age. Epidemiology is at present a hidden variable in many ecoimmunological studies; we should bring it to the light (Hawley and Altizer 2011).

Empirical studies testing our results face several challenges but also present opportunities for advancing the field of disease ecology. One such challenge is characterizing epidemiological risk and how it varies with age. Longitudinal studies may be our most promising avenue forward. For example, Combrink et al. (2020) monitored a herd of Cape buffalo across several years and discovered that some parasites were unlikely to be found in young buffalo. This suggests that infection risk may increase somewhat with age in this system. Force-of-infection models (e.g. Heisey et al. 2006, Pepin et al. 2017) may be particularly useful for establishing infection risk, which is highly influential in our analysis. An additional aspect of the challenge of ascertaining risk is that the variation within a population in epidemiological risk is important for assessing the amount of resolution we can obtain in comparative studies. The greater the variation between individuals in epidemiological risk, the greater the within-population variation in immune strategy, which in turn would spill over to reduced differences among populations in the distributions of optimal immune strategies each contains.

Another challenge for empirically testing our results is the development of methods to characterize immune sensitivity and/or specificity. One option is the quantity of cytokines or the dose-response curves of immune defenses produced in response to antigen exposure or experimental infection (e.g. Vedder et al. 1999, VanDalen et al. 2013) – these might offer a window into specificity and sensitivity as they shape what quantity of antigen is required to stimulate an immune response. This would potentially be straightforward, although changes in gene regulation might shape sensitivity and specificity such that they are dynamic properties across life (a possibility explored in Metcalf et al. 2017). Ascertaining incidence of immunopathology may also provide insight into immune strategy, because ultimately immunopathology should be more common in more sensitive organisms. Alternatively, to understand receptor specificity and the range of antigens provoking an immune response, one could directly examine binding affinities for TLRs or the specificity of B and T cell receptors and the activation thresholds for cells thus bound (Metcalf et al. 2017). Any of these approaches would help us identify the approach of a given organism’s immune system to identifying and responding to infections.

Lastly, while we use a demographic framework based on single-year age classes, and while we only focus on chordates, it is quite plausible that our results may apply more broadly to many different organisms with lifespans on many different temporal scales. For example, do tree seedlings in the undergrowth experience different risks of infection and mortality than adults stretching into the canopy? How do the temporal durations of these different growth stages affect overall disease hazard? Considering these two factors might help us gain insight into the immune defenses deployed by plants and other organisms. Our results might also offer insight into the immune strategies of microbial organisms. Past research has described how epidemiology can shape prokaryote immune defenses (Westra et al. 2015, Dimitriu et al. 2020, Chen et al. 2021), and our results may help us understand how risk variation and host strategy contribute. The general principle that we find here is that immune defense in chordates should be shaped according to how challenges differ across ages, and there is no reason that this would not apply across a wider range of host taxa.

We have quantified an important ecological interaction shaping optimal organismal immune strategy. In so doing, we show that differences in magnitude and timing of reproductive output can drive differences in immune strategy, and that different longevity-associated life history traits can pull in different directions on immune strategy. But epidemiology sets the strength and direction of the life history-immune strategy relationship for all life history traits.

## Materials and Methods

### Basic Model

We use a model of immune system recognition and response strategy first developed in Metcalf et al. (2017). In this model, immune strategy is described as a trade-off between sensitivity and specificity in detection and discernment of parasite signals (Fig. 1, Metcalf et al. 2017). The trade-off between sensitivity and specificity can be expressed mathematically with a receiver-operator curve (ROC), a tool frequently used in describing signal detection (Nesse 2005, Metcalf et al. 2017). Sensitivity and specificity are related with the equation

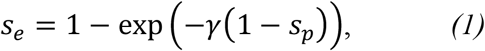

where *s_e_* is immune sensitivity, *s_p_* is immune specificity, and γ is a discrimination coefficient influenced by the overlap of the distributions of parasite and non-parasite signals and the ability of the host to discern that overlap (Fig. 1B, C). Higher values of γ indicate less overlap and greater discernment. Both *s_e_* and *s_p_* are constrained on the interval [0,1].

Overall mortality risk is a sum of mortality hazards from disease and non-disease causes. We include background, infection, and immunopathology mortality. Sensitivity and specificity, coupled with infection risk, influence the total mortality attributable to infection and immunopathology. Survival *s_x_* at age *x* is expressed via the following equation:

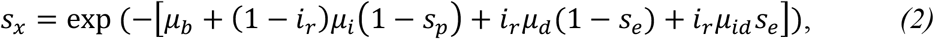

where *i_r_* is infection risk, *μ_b_* is background mortality risk, *μ_i_* is the risk of immunopathology mortality when uninfected, *μ_d_* is the risk of infection-induced mortality, assuming no immune response, and *μ_id_* is the risk of infection-induced mortality when there is an immune response, including any immunopathology risk. While the various mortality parameters are constrained only to be 0 or positive, *i_r_* is constrained on [0,1]. Substituting Equation (1) into Equation (2) allows calculation of *s_x_* as a function of *s_p_*.

When we discuss epidemiological risk in this paper, we are referring to any of *i_r_*, *μ_i_*, *μ_d_*, and *μ_id_*. By varying these parameters with age we can explore how immune strategy affects survival at different ages when the risks differ during life. We combine the resulting survival curve with a reproductive output schedule to produce a demographic matrix with survival and fertility parameters for multiple age classes. This matrix describes the dynamics of a population with the given background mortality, epidemiological risks, reproductive output schedule, and immune strategy. As in Metcalf and Graham (2018), we define the fitness of a population as λ, the dominant eigenvalue of the matrix, which is the population growth rate (Caswell 2001); this incorporates a temporal component that generally makes it a preferable fitness measure to R_0_, the net reproductive rate (Metcalf and Pavard 2007). By calculating λ for a range of immune strategies – here *s_p_* values from 0 to 1 with intervals of 0.001 and the associated values of *s_e_* – we can identify the strategy maximizing fitness. While we mostly report our results as optimal immune specificity for convenience, all values can be transformed to optimal immune sensitivities by Equation (1).

### Designating Reproductive and Epidemiological Variation

In our exploration of the effects of reproductive demography, we defined five different schedules for reproductive output. For our baseline strategy, reproduction was constant at all ages from the age of first reproduction (“Baseline”). We then chose two strategies that also featured constant reproduction across age, but with higher (“High”) or lower (“Low”) yearly reproductive outputs relative to that baseline. To investigate how variation in reproduction with age might affect optimal immune strategy, we also used simple schedules where reproduction either increased with age at a constant rate from a low initial value (“Rising”) or decreased with age at a constant rate from a high initial value (“Declining”). For each of these reproductive schedules we calculated the optimal immune strategy, as described above, in a particular epidemiological context where infection risk drops at the age of reproductive maturity. In addition, changing reproductive output alters fitness, such that our five schedules have different fitness values when confronted with the same mortality risks; therefore, to establish the robusticity of our results, we also explored an alternate case where background mortality *μ_b_* was manipulated to approximately equalize the fitness values associated with each schedule while still using the same epidemiological context across the five schedules.

We next considered how variation in epidemiological risk across life affects optimal immune strategy. We identified two epidemiological scenarios, each with a single change in risk at reproductive maturity – our stepped mode (Table S1). For our main results we focused on variation in *i_r_*. In our scenarios *i_r_* fell (A1) and rose (A2) at reproductive maturity, respectively. For all scenarios, the other parameters remained constant. We then combined these two scenarios with the five different reproductive schedules to explore how varying epidemiological context alters the relationship between reproduction and optimal immune strategy. As previously shown, if epidemiological risk does not vary with age, features of demography do not modulate the optimal immune strategy (Metcalf and Graham 2018) and therefore, we do not further evaluate scenarios where epidemiological risk is flat over age.

We also explored how different magnitudes of risk, and quantities of variation in said risk, affected optimal immune strategy. Here we explored four different scenarios for variation in *i_r_*, again with risk changing once at reproductive maturity (Table S2). To explore the importance of absolute magnitude of risk, we used two scenarios where *i_r_* declined across life; the amount of decline was the same between the two scenarios, but the absolute level of risk differed in each age class, with a low-risk scenario (B1) and a high-risk scenario (B2). To explore how magnitude of risk variation affected optimal strategy, we used two scenarios with different quantities of risk change at reproductive maturity, with a large-change scenario (B3) and a small-change scenario (B4). For all scenarios, the other parameter values in our model were *μ_b_* = 0.15, *μ_d_* = 0.3, *μ_i_* = 0.1, *μ_id_* = 0.01, and γ = 4. As above, we combined these scenarios with the five reproductive schedules to investigate how magnitude and variation of risk affect optimal immune strategies.

To further consider the importance of which risk parameters vary with age and how they do so, we also considered two alternative types of variation in risk with age. For the first type, we varied *μ_d_* with age while *i_r_* was held constant. We repeated each of the A and B scenarios with variation in *μ_d_*. In these scenarios, *i_r_* = 0.4 at all ages, while *μ_d_* varies on the same intervals that *i_r_* did in the analogous scenarios, and all other parameters retain the same values. For the second type, we defined a separate mode of variation – “smoothed” – in which risk change between age classes is constant. To translate our stepped scenarios to our smoothed scenarios, we used the risk value prior to reproduction as the risk value for the first age class, and we used the risk value for reproductive years as the risk value for the last age class. We used this approach with both *i_r_* and *μ_d_*, with the same parameter values. We therefore explored versions of A1–A2 and B1–B4 for both the stepped and smoothed modes and for both *i_r_* and *μ_d_*.

### Predicting Optimal Immune Defense from Population Projection Matrices

To root our analysis in existing schedules of reproduction and survival, we leveraged a large representative dataset of population projection matrices from the COMADRE database (Salguero-Gómez et al. 2016, COMADRE Animal Matrix Database 2020) to serve as starting points for the analysis. These matrices give us survival and fertility parameters measured from wild populations of various animal species. We combined these parameters, with some transformation, with epidemiological risk parameters and then used our range of *s_p_* and *s_e_* values to determine what immune strategy would produce the maximum population growth rate given those risk, fertility, and survival parameters.

We restricted our analysis to primitive, irreducible, and ergodic matrices (i.e., the matrix diagonal contains 0s except for the final entry, survival is contained in the off-diagonals, and fertility is contained in the first row) that are structured by age in years and for which each age class is equivalent to one year. By using only matrices structured by years we ensured comparability between populations in epidemiology and life history. After filtering, we were left with 298 population matrices representing 129 chordate species. We also report results for an analysis that includes only 151 population matrices representing 47 mammal species.

### Transforming Matrices for Use with Prediction Method

Population projection matrices can be of varying size, depending on both the demography of the population and the data available. For matrices with each age class equivalent to one year, which we are using, this primarily affects the amount of detail in describing the population dynamics of later age classes. Because we are interested in how epidemiological risk variation across life affects immune strategy, we standardized matrix dimension with respect to checkpoints in the life history of an organism. We transformed each original matrix to a new one with its dimension the sum of the age at first reproduction of the organism and the reproductive life expectancy of the organism. In this way we can scale epidemiological risk variation over age to each species’s lifespan while retaining comparability across taxa with respect to life history.

Re-sizing matrices according to our above criterion requires increasing the dimension of some matrices and decreasing the dimension of others. Increasing dimension is simple: we added columns and rows with the same survival and fertility parameters as those in the final column and row of the original matrix. Such a matrix describes the same population dynamics, including λ, as the original matrix, because the final age class of any matrix describes the survival and fertility of any individual of that age or older. Decreasing dimension is a slightly more complex process. We used an algorithm designed by Hooley (2000). In brief, this algorithm collapses designated rows and columns by taking an average of the survival and fertility parameters in those rows and columns and weighting by stable age structure in those age classes. The weighted average parameters are used as the fertility and survival parameters for a single new column replacing the collapsed columns. We follow the recommendation from Salguero-Gómez and Plotkin (2010) to collapse the rows and columns that represent the oldest age classes, as this criterion minimizes the distortion of demographic patterns by only affecting the degree of nuance in later age classes, which will generally have relatively small contributions to fitness. This collapsing method produces a matrix of the desired dimension with population dynamics quite tightly approximating those of the original matrix. Overall 288 of the 298 matrices in our dataset required resizing, but this resizing has minimal impact on estimates of population dynamics for each matrix (Fig. S1).

While the COMADRE matrices do provide the age trajectory of survival (and thus mortality hazard) for our focal species, they offer no insight into causes of mortality. In theory one would want to determine immune strategy based on a precise knowledge of infection and immunopathology-associated risk. But there is no way to establish what fraction of the mortality recorded in the matrix is attributable to infectious disease or immunopathology vs. other causes of death. To address this, we must define our own background mortality and epidemiological risk parameters, and these can only be loose approximations of the risks experienced by a given organism as recorded in the relevant population projection matrix. Because it has previously been established that epidemiological context strongly affects immune strategy in this model framework (Metcalf et al. 2017, Metcalf and Graham 2018), we want to compare among species subject to the same epidemiological risks to understand how life history might be linked to immune strategy without confounding. We therefore choose arbitrary epidemiological risk parameters (and schedules of variation thereof) which can be defined consistently for all matrices.

Our remaining problem is choosing background mortality parameter values. Because we are deriving life history from the original population projection matrix, we want to choose our value for *μ_b_* based on the mortality recorded in the original matrix. We therefore designate *μ_b_* for each age class of our new matrix as being some proportion *ρ* of the original mortality in that age class in the original matrix. This designation arbitrarily defines *ρ* as the proportion of overall mortality attributable to non-disease causes, while 1 – *ρ* is the proportion associated with disease and immunopathology. Thus the *μ_b_* value for a given age *x* in our new survival curve is calculated as a product of the survival *s_x_* in the relevant age class *x* of the re-sized matrix and a constant *ρ*, such that

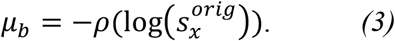

For our analysis we used *ρ* = 0.75, but the value of *ρ* has a minimal effect on our results. We combine the new *μ_b_* value for each age class calculated as in Equation (3) with epidemiological risks to produce a new survival curve for the population under consideration. The resulting survival curve only approximates the original survival curve, but it allows us to investigate different epidemiological scenarios and immune strategies.

Because the qualitative results from variation in *i_r_* and *μ_d_* are the same in our earlier analyses, we only looked at different *i_r_* scenarios for this portion of our analysis. We considered two ways that *i_r_* might vary with age. The first is identical to our stepped mode described earlier: we define one *i_r_* value for pre-reproductive age classes and another *i_r_* value for reproductive age classes. Our second mode is similar to our smoothed mode previously described in allowing *i_r_* to change smoothly across lifespan. Here, we define a starting risk for the schedule, which would be *i_r_* in the first age class of the matrix, and an ending risk for the schedule, which would be *i_r_* for the last age class of the matrix, and allow a uniform change in *i_r_* across intermediate age classes. However, since each matrix differs in dimension (and therefore absolute time and age described), this rate of change in *i_r_* cannot be the same from matrix to matrix (Fig. S2A). Therefore we scale the rate of change by the dimension of the matrix, such that change in *i_r_* relative to the demography described by the matrix – demography that is standardized as the sum of age at first reproduction and reproductive life expectancy – is constant across all populations. An example of how this works: in a matrix with dimension 3 a schedule for *i_r_* might be [0.45, 0.325, 0.2], while for a matrix of dimension 6 the equivalent schedule for *i_r_* would be [0.45, 0.4, 0.35, 0.3, 0.25, 0.2]. As noted above, the absolute pace of change of risk varies between the two schedules, but the relative pace of change with respect to life history is constant and the absolute change across life is also constant (Fig. S2B).

In this way we produce equations for *s_x_* for each age class in the matrix. With these, and the fertility parameters of the original matrix, we determined optimal specificity as described above, creating matrices across the range of values of *s_p_* and identifying the values of *s_p_* and *s_e_* that produce λ_max_. This is our predicted optimal immune strategy associated with the original matrix from the COMADRE database. As above, we present our results in terms of specificity, but *s_p_* values can be easily translated to *s_e_* values by Equation 1.

### Linear Model for Relationship between Demography and Predicted Immune Strategy

To describe the relationship between life history and immune strategy, we calculated four life history summary statistics from our matrices: age class of first reproduction (AFR), life expectancy post-reproductive maturity or reproductive life expectancy (RLE), mean reproductive rate (MRR), and generation time (GT). Each of these statistics is calculated in the manner defined by Healy et al. (2019). To describe the relationship between the life history traits and the predicted optimal immune specificity, we used a Bayesian linear model of specificity as a function of life history trait values. For our main results we used life history statistics as calculated from the original matrix in the COMADRE database, rather than the transformed, optimized matrix, to counteract any systematic distortions introduced by our manipulation of matrix dimension and survival curve calculation (although note as above that dimension manipulation does not seem to induce any demographic distortions). However, to ensure robusticity we also did repeat our analysis using statistics calculated from the post-processing matrices. In addition, we considered GT separately from the other three traits because it is highly collinear with each of them in our dataset.

Life history trait parameter values were first log-transformed to produce trait value distributions approximating a normal distribution. They were then standardized by calculating Z-scores from the mean and standard deviation of the sampled values for each parameter. Predicted optimal immune sensitivity *s_p_*^*^ was left unstandardized because it has no units and is constrained on the interval [0,1]. We also assumed for our linear model that *s_p_*^*^ is normally distributed; while *s_p_* does have a constrained possible range, our model predictions never approached those boundaries, such that a normal distribution can reasonably be used. The model was constructed with the “ulam” function of the “rethinking” package v2.01 (which implements RStan) in R v4.0.1 and run for 2000 samples (McElreath 2020, R Core Team 2020). We designated the prior for the intercept of *s_p_*^*^ as a normal distribution with μ = 0.5 and σ = 0.15; for all coefficients, we similarly used a normal distribution prior centered on μ = 0, with σ = 0.1. In total we examined 121 different scenarios for variation in infection risk, with differences in how much infection risk changed across life and whether it rose or fell. We produced predicted optimal immune specificities for our set of population projection matrices, and we analyzed the results with linear models for each scenario. We repeated these analyses for both stepped and smoothed modes of *i_r_* variation.

## Supporting information

R code used in analyses

## Acknowledgments

We thank C. Riehl, B. vonHoldt, and the Graham Lab for valuable discussions. A. E. D. acknowledges funding support from the US National Science Foundation Graduate Research Fellowship Program (Award Number DGE-2039656). A. M. acknowledges funding support from a Princeton University Lewis-Sigler Fellowship.

## Data and Materials Availability

The R code used in this paper is available as a supplemental file with this manuscript. The COMADRE database of animal life history matrices is available at https://compadre-db.org/Data/Comadre.

## References

Albery, G. F., & Becker, D. J. (2021). Fast-lived Hosts and Zoonotic Risk. Trends in Parasitology, 37(2), 117–129. https://doi.org/10.1016/j.pt.2020.10.012

Altizer, S., Nunn, C. L., Thrall, P. H., Gittleman, J. L., Antonovics, J., Cunningham, A. A., Dobson, A. P., Ezenwa, V., Jones, K. E., Pedersen, A. B., Poss, M., & Pulliam, J. R. C. (2003). Social Organization and Parasite Risk in Mammals: Integrating Theory and Empirical Studies. Annual Review of Ecology, Evolution, and Systematics, 34(1), 517–547. https://doi.org/10.1146/annurev.ecolsys.34.030102.151725

Antonovics, J., Boots, M., Abbate, J., Baker, C., McFrederick, Q., & Panjeti, V. (2011). Biology and evolution of sexual transmission. Annals of the New York Academy of Sciences, 1230(1), 12–24. https://doi.org/10.1111/j.1749-6632.2011.06127.x

Best, A., & Hoyle, A. (2013). The evolution of costly acquired immune memory. Ecology and Evolution, 3(7), 2223–2232. https://doi.org/10.1002/ece3.611

Boots, M., Donnelly, R., & White, A. (2013). Optimal immune defence in the light of variation in lifespan. Parasite Immunology, 35(11), 331–338. https://doi.org/10.1111/pim.12055

Boots, Michael, Best, A., Miller, M. R., & White, A. (2009). The role of ecological feedbacks in the evolution of host defence: What does theory tell us? Philosophical Transactions of the Royal Society B: Biological Sciences, 364(1513), 27–36. https://doi.org/10.1098/rstb.2008.0160

Caswell, H. (2001). Matrix Population Models: Construction, Analysis, and Interpretation (Second Edition). Sinauer Associates, Inc.

Chen, H., Mayer, A., & Balasubramanian, V. (2021). A scaling law in CRISPR repertoire sizes arises from avoidance of autoimmunity. BioRxiv, 2021.01.04.425308. https://doi.org/10.1101/2021.01.04.425308

COMADRE Animal Matrix Database (2021). Available from https://www.compadre-db.org [31 January 2021, 4.21.1.0].

Combrink, L., Glidden, C. K., Beechler, B. R., Charleston, B., Koehler, A. V., Sisson, D., Gasser, R. B., Jabbar, A., & Jolles, A. E. (2020). Age of first infection across a range of parasite taxa in a wild mammalian population. Biology Letters, 16(2), 20190811. https://doi.org/10.1098/rsbl.2019.0811

Cressler, C. E., Graham, A. L., & Day, T. (2015). Evolution of hosts paying manifold costs of defence. Proceedings of the Royal Society B: Biological Sciences, 282(1804), 20150065. https://doi.org/10.1098/rspb.2015.0065

Dimitriu, T., Szczelkun, M. D., & Westra, E. R. (2020). Evolutionary Ecology and Interplay of Prokaryotic Innate and Adaptive Immune Systems. Current Biology, 30(19), R1189–R1202. https://doi.org/10.1016/j.cub.2020.08.028

Donnelly, R., White, A., & Boots, M. (2015). The epidemiological feedbacks critical to the evolution of host immunity. Journal of Evolutionary Biology, 28(11), 2042–2053. https://doi.org/10.1111/jeb.12719

Donnelly, R., White, A., & Boots, M. (2017). Host lifespan and the evolution of resistance to multiple parasites. Journal of Evolutionary Biology, 30(3), 561–570. https://doi.org/10.1111/jeb.13025

Downs, C. J., Dochtermann, N. A., Ball, R., Klasing, K. C., & Martin, L. B. (2020). The Effects of Body Mass on Immune Cell Concentrations of Mammals. The American Naturalist, 195(1), 107–114. https://doi.org/10.1086/706235

Duffy, M. A., Ochs, J. H., Penczykowski, R. M., Civitello, D. J., Klausmeier, C. A., & Hall, S. R. (2012). Ecological Context Influences Epidemic Size and Parasite-Driven Evolution. Science, 335(6076), 1636–1638. https://doi.org/10.1126/science.1215429

Graham, A. L., Allen, J. E., & Read, A. F. (2005). Evolutionary Causes and Consequences of Immunopathology. Annual Review of Ecology, Evolution, and Systematics, 36(1), 373–397. https://doi.org/10.1146/annurev.ecolsys.36.102003.152622

Hamilton, R., Siva-Jothy, M., & Boots, M. (2008). Two arms are better than one: Parasite variation leads to combined inducible and constitutive innate immune responses. Proceedings of the Royal Society B: Biological Sciences, 275(1637), 937–945. https://doi.org/10.1098/rspb.2007.1574

Hawley, D. M., & Altizer, S. M. (2011). Disease ecology meets ecological immunology: Understanding the links between organismal immunity and infection dynamics in natural populations. Functional Ecology, 25(1), 48–60. https://doi.org/10.1111/j.1365-2435.2010.01753.x

Healy, K., Ezard, T. H. G., Jones, O. R., Salguero-Gómez, R., & Buckley, Y. M. (2019). Animal life history is shaped by the pace of life and the distribution of age-specific mortality and reproduction. Nature Ecology & Evolution, 3(8), 1217–1224. https://doi.org/10.1038/s41559-019-0938-7

Heisey, D. M., Joly, D. O., & Messier, F. (2006). The Fitting of General Force-of-Infection Models to Wildlife Disease Prevalence Data. Ecology, 87(9), 2356–2365. https://doi.org/10.1890/0012-9658(2006)87[2356:TFOGFM]2.0.CO;2

Hooley, D. E. (2000). Collapsed Matrices with (Almost) the Same Eigenstuff. The College Mathematics Journal, 31(4), 297–299. https://doi.org/10.2307/2687420

Hudson, P. J., & Dobson, A. P. (1995). Macroparasites: Observing patterns in naturally fluctuating animal populations. In B. T. Grenfell & A. P. Dobson (Eds.), Ecology of Infectious Diseases in Natural Populations (Vol. 7) (pp. 144–176). Cambridge University Press.

Knowles, S. C. L., Nakagawa, S., & Sheldon, B. C. (2009). Elevated reproductive effort increases blood parasitaemia and decreases immune function in birds: A meta-regression approach. Functional Ecology, 23(2), 405–415. https://doi.org/10.1111/j.1365-2435.2008.01507.x

Kubinak, J. L., Ruff, J. S., Hyzer, C. W., Slev, P. R., & Potts, W. K. (2012). Experimental viral evolution to specific host MHC genotypes reveals fitness and virulence trade-offs in alternative MHC types. Proceedings of the National Academy of Sciences, 109(9), 3422–3427. https://doi.org/10.1073/pnas.1112633109

Lazzaro, B. P., & Little, T. J. (2009). Immunity in a variable world. Philosophical Transactions of the Royal Society B: Biological Sciences, 364(1513), 15–26. https://doi.org/10.1098/rstb.2008.0141

Lee, K. A., Wikelski, M., Robinson, W. D., Robinson, T. R., & Klasing, K. C. (2008). Constitutive immune defences correlate with life-history variables in tropical birds. Journal of Animal Ecology, 77(2), 356–363. https://doi.org/10.1111/j.1365-2656.2007.01347.x

Lee, K. A. (2006). Linking immune defenses and life history at the levels of the individual and the species. Integrative and Comparative Biology, 46(6), 1000–1015. https://doi.org/10.1093/icb/icl049

Lochmiller, R. L., & Deerenberg, C. (2000). Trade-offs in evolutionary immunology: Just what is the cost of immunity? Oikos, 88(1), 87–98. https://doi.org/10.1034/j.1600-0706.2000.880110.x

Martin, L. B., Weil, Z. M., & Nelson, R. J. (2007). Immune Defense and Reproductive Pace of Life in Peromyscus Mice. Ecology, 88(10), 2516–2528. https://doi.org/10.1890/07-0060.1

Mayer, A., Balasubramanian, V., Mora, T., & Walczak, A. M. (2015). How a well-adapted immune system is organized. Proceedings of the National Academy of Sciences, 112(19), 5950–5955. https://doi.org/10.1073/pnas.1421827112

Mayer, A., Mora, T., Rivoire, O., & Walczak, A. M. (2016). Diversity of immune strategies explained by adaptation to pathogen statistics. Proceedings of the National Academy of Sciences, 113(31), 8630–8635. https://doi.org/10.1073/pnas.1600663113

McElreath, R. (2020). Rethinking: Statistical Rethinking book package. R package v. 2.01. Available from https://github.com/rmcelreath/rethinking

Medzhitov, R., Schneider, D. S., & Soares, M. P. (2012). Disease Tolerance as a Defense Strategy. Science, 335(6071), 936–941. https://doi.org/10.1126/science.1214935

Metcalf, C. J. E., & Graham, A. L. (2018). Schedule and magnitude of reproductive investment under immune trade-offs explains sex differences in immunity. Nature Communications, 9(1), 1–9. https://doi.org/10.1038/s41467-018-06793-y

Metcalf, C. J. E., & Pavard, S. (2007). Why evolutionary biologists should be demographers. Trends in Ecology & Evolution, 22(4), 205–212. https://doi.org/10.1016/j.tree.2006.12.001

Metcalf, C. J. E., Tate, A. T., & Graham, A. L. (2017). Demographically framing trade-offs between sensitivity and specificity illuminates selection on immunity. Nature Ecology & Evolution, 1(11), 1766–1772. https://doi.org/10.1038/s41559-017-0315-3

Miller, M. R., White, A., & Boots, M. (2007). Host Life Span and the Evolution of Resistance Characteristics. Evolution, 61(1), 2–14. https://doi.org/10.1111/j.1558-5646.2007.00001.x

Nesse, R. M. (2005). Natural selection and the regulation of defenses: A signal detection analysis of the smoke detector principle. Evolution and Human Behavior, 26(1), 88–105. https://doi.org/10.1016/j.evolhumbehav.2004.08.002

Norris, K., & Evans, M. R. (2000). Ecological immunology: Life history trade-offs and immune defense in birds. Behavioral Ecology, 11(1), 19–26. https://doi.org/10.1093/beheco/11.1.19

Nunn, C. L. (2002). A Comparative Study of Leukocyte Counts and Disease Risk in Primates. Evolution, 56(1), 177–190. https://doi.org/10.1111/j.0014-3820.2002.tb00859.x

Pepin, K. M., Kay, S. L., Golas, B. D., Shriner, S. S., Gilbert, A. T., Miller, R. S., Graham, A. L., Riley, S., Cross, P. C., Samuel, M. D., Hooten, M. B., Hoeting, J. A., Lloyd-Smith, J. O., Webb, C. T., & Buhnerkempe, M. G. (2017). Inferring infection hazard in wildlife populations by linking data across individual and population scales. Ecology Letters, 20(3), 275–292. https://doi.org/10.1111/ele.12732

R Core Team. (2020). R: A language and environment for statistical computing. Vienna, Austria. Retrieved from https://www.cran.r-project.org/

Råberg, L., Graham, A. L., & Read, A. F. (2009). Decomposing health: Tolerance and resistance to parasites in animals. Philosophical Transactions of the Royal Society B: Biological Sciences, 364(1513), 37–49. https://doi.org/10.1098/rstb.2008.0184

Ricklefs, R. E., & Wikelski, M. (2002). The physiology/life-history nexus. Trends in Ecology & Evolution, 17(10), 462–468. https://doi.org/10.1016/S0169-5347(02)02578-8

Ronget, V., & Gaillard, J.-M. (2020). Assessing ageing patterns for comparative analyses of mortality curves: Going beyond the use of maximum longevity. Functional Ecology, 34(1), 65–75. https://doi.org/10.1111/1365-2435.13474

Salguero-Gómez, R., Jones, O. R., Archer, C. R., Bein, C., Buhr, H. de, Farack, C., Gottschalk, F., Hartmann, A., Henning, A., Hoppe, G., Römer, G., Ruoff, T., Sommer, V., Wille, J., Voigt, J., Zeh, S., Vieregg, D., Buckley, Y. M., Che-Castaldo, J.,… Vaupel, J. W. (2016). COMADRE: A global data base of animal demography. Journal of Animal Ecology, 85(2), 371–384. https://doi.org/10.1111/1365-2656.12482

Salguero-Gómez, R., & Plotkin, J. B. (2010). Matrix Dimensions Bias Demographic Inferences: Implications for Comparative Plant Demography. The American Naturalist, 176(6), 710–722. https://doi.org/10.1086/657044

Schmid-Hempel, P. (2003). Variation in immune defence as a question of evolutionary ecology. Proceedings of the Royal Society of London. Series B: Biological Sciences, 270(1513), 357–366. https://doi.org/10.1098/rspb.2002.2265

Sheldon, B. C., & Verhulst, S. (1996). Ecological immunology: Costly parasite defences and trade-offs in evolutionary ecology. Trends in Ecology & Evolution, 11(8), 317–321. https://doi.org/10.1016/0169-5347(96)10039-2

Shudo, E., & Iwasa, Y. (2001). Inducible Defense against Pathogens and Parasites: Optimal Choice among Multiple Options. Journal of Theoretical Biology, 209(2), 233–247. https://doi.org/10.1006/jtbi.2000.2259

Silk, M. J., & Hodgson, D. J. (2021). Life history and population regulation shape demographic competence and influence the maintenance of endemic disease. Nature Ecology & Evolution, 5(1), 82–91. https://doi.org/10.1038/s41559-020-01333-8

Stearns, S. C. (1992). The Evolution of Life Histories. Oxford University Press.

Tella, J. L., Scheuerlein, A., & Ricklefs, R. E. (2002). Is cell–mediated immunity related to the evolution of life-history strategies in birds? Proceedings of the Royal Society of London. Series B: Biological Sciences, 269(1495), 1059–1066. https://doi.org/10.1098/rspb.2001.1951

Valenzuela-Sánchez, A., Wilber, M. Q., Canessa, S., Bacigalupe, L. D., Muths, E., Schmidt, B. R., Cunningham, A. A., Ozgul, A., Johnson, P. T. J., & Cayuela, H. (2021). Why disease ecology needs life-history theory: A host perspective. Ecology Letters, *n/a*(n/a). https://doi.org/10.1111/ele.13681

van Boven, M., & Weissing, F. J. (2004). The Evolutionary Economics of Immunity. The American Naturalist, 163(2), 277–294. https://doi.org/10.1086/381407

VanDalen, K. K., Hall, J. S., Clark, L., McLean, R. G., & Smeraski, C. (2013). West Nile Virus Infection in American Robins: New Insights on Dose Response. PLOS ONE, 8(7), e68537. https://doi.org/10.1371/journal.pone.0068537

Vedder, H., Schreiber, W., Yassouridis, A., Gudewill, S., Galanos, C., & Pollmächer, T. (1999). Dose-dependence of bacterial lipopolysaccharide (LPS) effects on peak response and time course of the immune-endocrine host response in humans. Inflammation Research, 48(2), 67–74. https://doi.org/10.1007/s000110050408

Westra, E. R., van Houte, S., Oyesiku-Blakemore, S., Makin, B., Broniewski, J. M., Best, A., Bondy-Denomy, J., Davidson, A., Boots, M., & Buckling, A. (2015). Parasite Exposure Drives Selective Evolution of Constitutive versus Inducible Defense. Current Biology, 25(8), 1043–1049. https://doi.org/10.1016/j.cub.2015.01.065

Wilson, K., Bjørnstad, O. N., Dobson, A. P., Merler, S., Poglayen, G., Randolph, S. E., Read, A. F., Skorping, A. (2002). Heterogeneities in microparasite infections: patterns and processes. In P. J. Hudson, A. P. Rizzoli, B. T. Grenfell, J. A. P. Heesterbeek, A. P. Dobson. Ecology of Wildlife Diseases (pp. 6–44). Oxford University Press.

Zuk, M., & Stoehr, A. M. (2002). Immune Defense and Host Life History. The American Naturalist, 160(S4), S9–S22. https://doi.org/10.1086/342131

