## Supplementary Materials for "Optimal immune specificity at the intersection of host life history and parasite epidemiology"

For the code used for our analyses and generating the figures, see the “Downieetal2021.R” file.

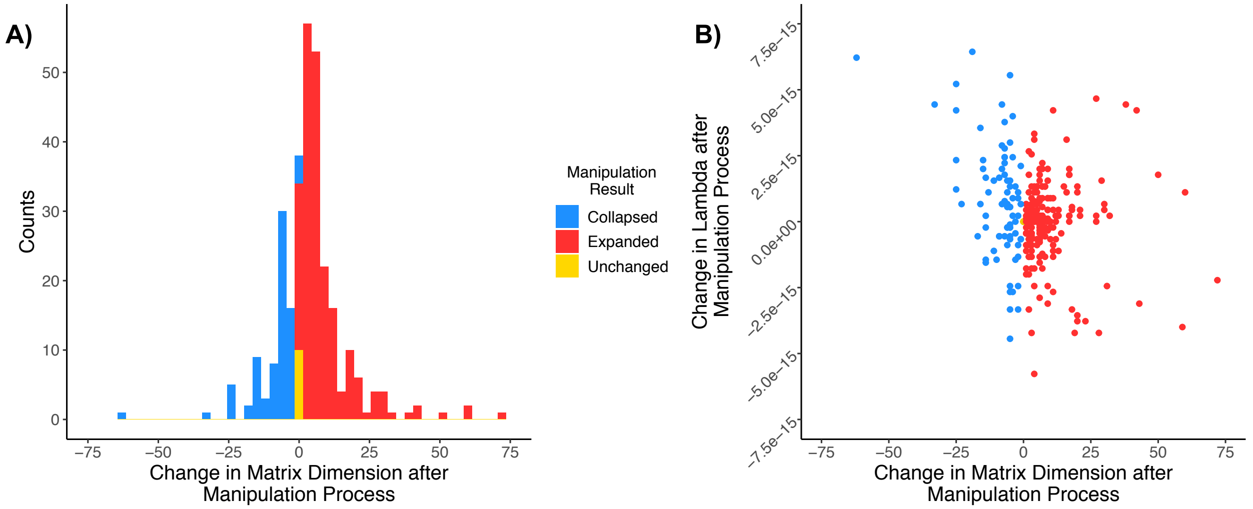


**Supplementary Figure 1: Implementation of matrix manipulation methods does not distort demographic patterns.** A) Histogram shows, for the 298 matrices used in the COMADRE method, how they were altered in dimension, either expanded or collapsed, and the change in dimension after alteration. B) Plot shows the change in matrix dimension and the associated difference between λ when calculated for the original matrix and when calculated for the matrix after the dimension has been altered.

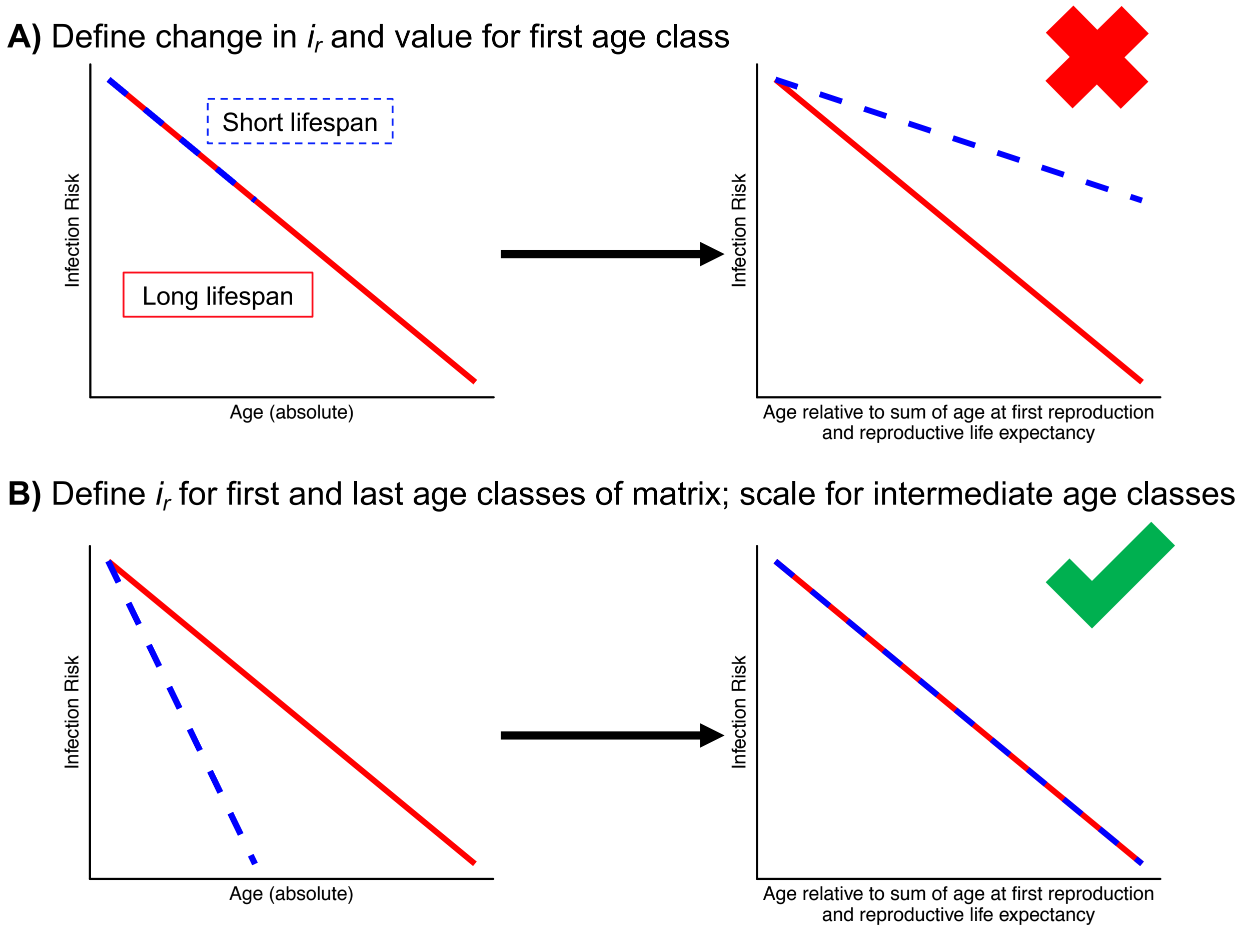


**Supplementary Figure 2: Method for scaling risk parameter change to matrix dimension.** X-coordinates for lines in absolute age plots determined by age classes within matrix, and so the blue line is for an organism with a short lifespan and smaller matrix dimension. Dimension of manipulated matrices (see Fig. S1) determined as sum of age at first reproduction and reproductive life expectancy. A) Picking a constant interval of change from one age class to another produces a schedule that is consistent neither in range of risk across life nor change in risk relative to defined aspects of life history. B) Defining start and end points for risk parameter ensures that range of risk across life is consistent, as is change in risk relative to life history.

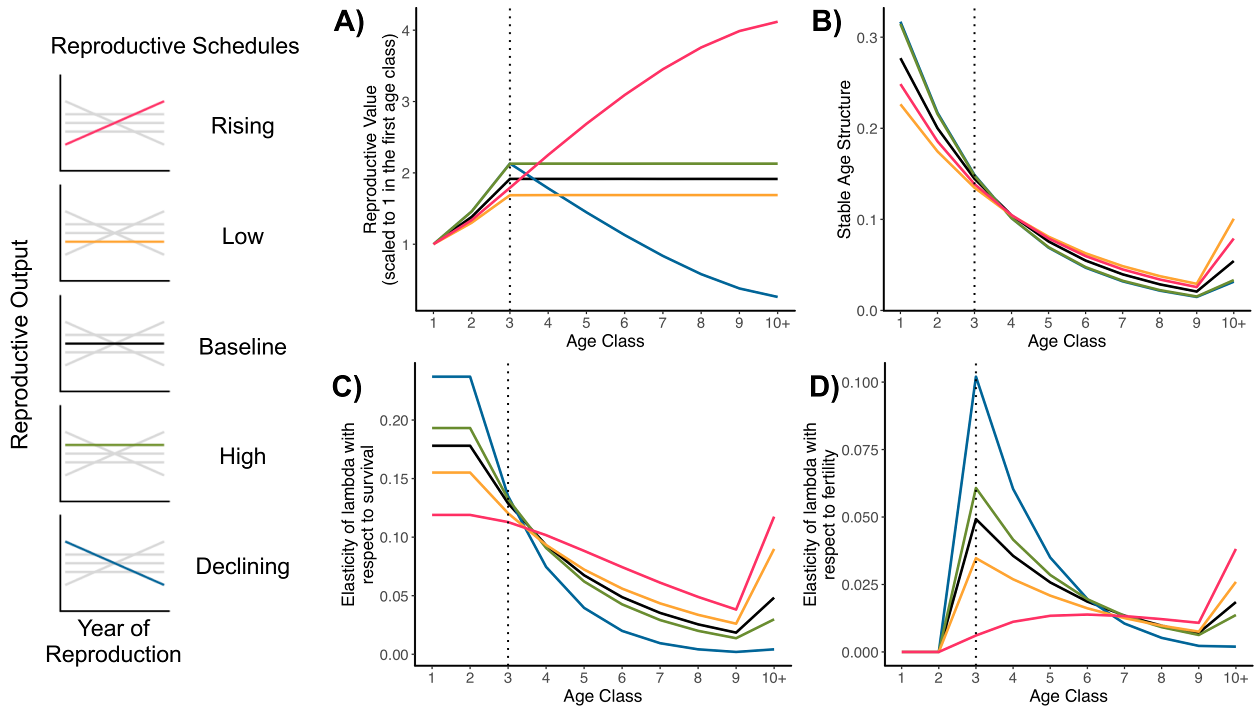


**Supplementary Figure 3: Demographic details for reproductive schedules with optimal immune strategies.** Optimal immune strategies identified in Figure 2 and Table S3. Dashed line shows age class of reproductive maturity, the third age class. A) Reproductive value distributions, with reproductive value in the first age class defined as 1; B) Stable age structures; C) Elasticities of $\lambda$ with respect to survival; D) Elasticities of $\lambda$ with respect to fertility. Infection risk $i_r$ declines from 0.6 in the first age class to 0.2 in the final age class. Other parameter values are $\mu_b = 0.15$, $\mu_i = 0.1$, $\mu_d = 0.3$, $\mu_{di} = 0.01$, and $\gamma = 4$.

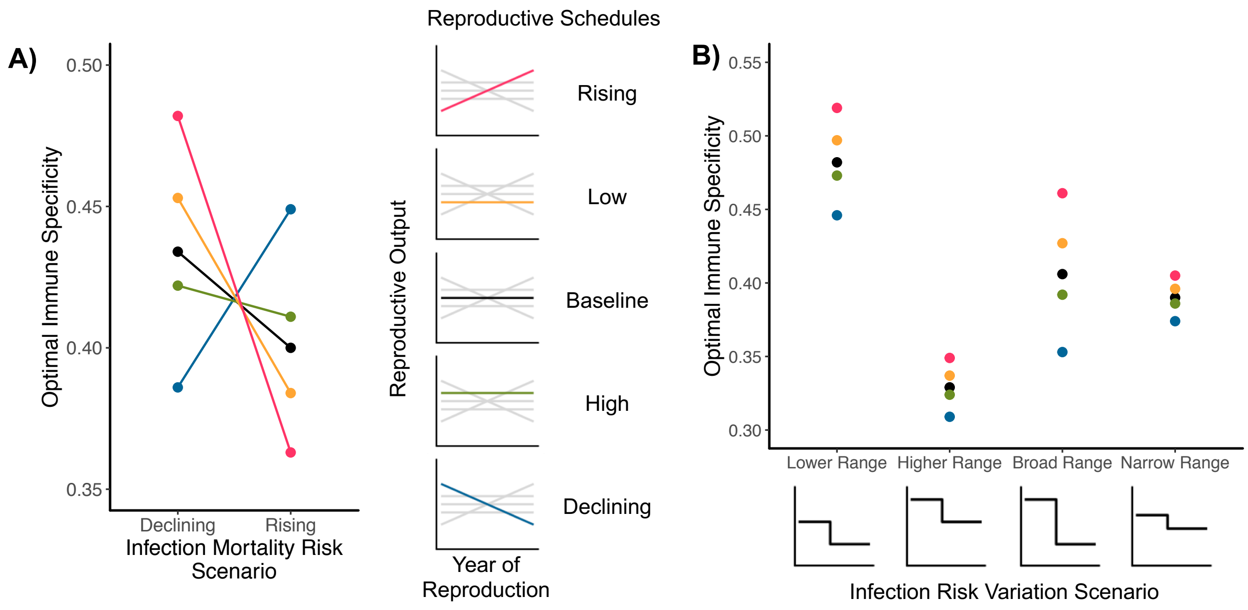


**Supplementary Figure 4: Epidemiological environment alters the effect of reproduction on immune strategy: infection mortality risk $\mu_d$.** Reproduction begins in the third age class for all schedules. In each scenario, $\mu_d$ starts at one value and drops to a lower value at the third age class. A) The change in optimal immune specificity associated with differences in epidemiological context (i.e. changes in $\mu_d$, on the x-axis) and reproduction (different points and lines). Parameter values are $\mu_b = 0.15$, $\mu_i = 0.1$, $\mu_{id} = 0.01$, $i_r = 0.4$, and $\gamma = 4$. In the declining scenario, $\mu_d$ declines with age from 0.6 to 0.2; in the rising scenario, $\mu_d$ increases from 0.2 to 0.6. B) The change in range of optimal specificities associated with different reproductive demographies associated with different magnitudes of variation in decline of infection mortality risk $\mu_d$ with age. Parameter values are $\mu_b = 0.15$, $\mu_i = 0.1$, $\mu_{id} = 0.01$, $i_r = 0.4$, and $\gamma = 4$. In the lower range scenario, $\mu_d$ declines from 0.45 to 0.2; in the higher range, from 0.7 to 0.45; in the broad range, from 0.7 to 0.2; in the narrow range, from 0.525 to 0.375.

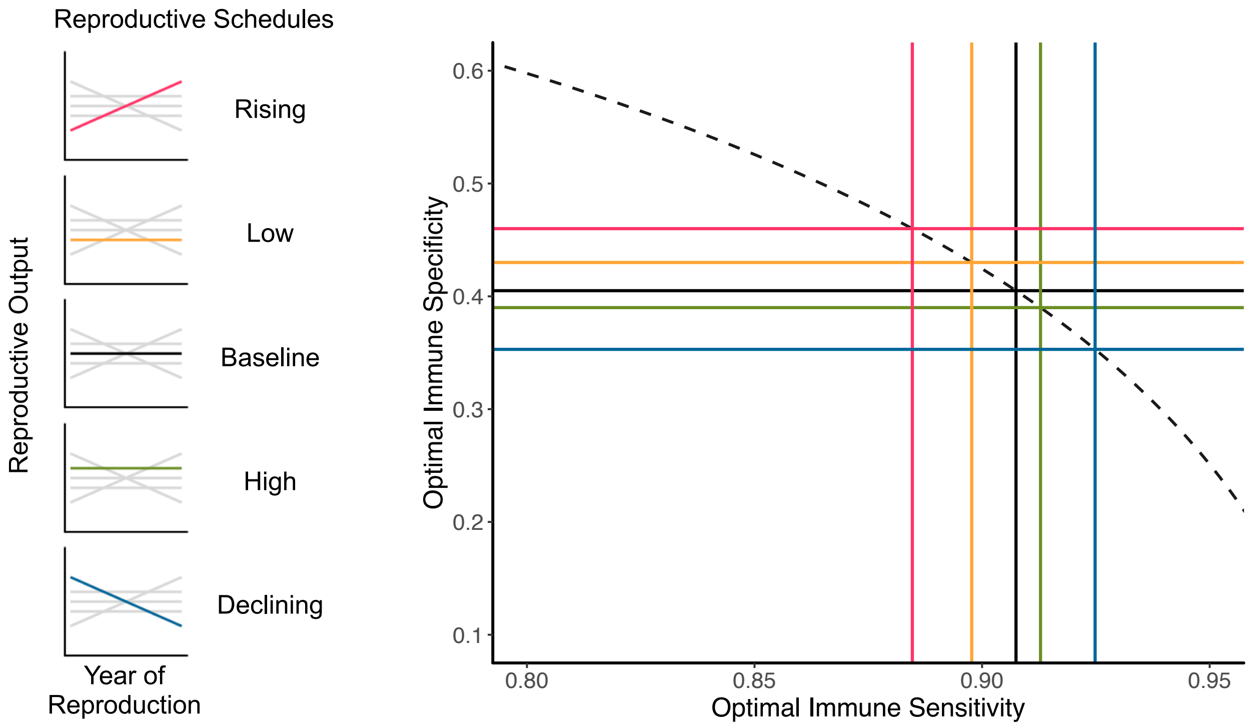


**Supplementary Figure 5: Influence of reproductive demography on optimal immune strategy: smoothed infection risk variation.** Plot showing optimal combination of immune sensitivity and specificity for each of five reproductive demographic schedules (at left), for a single epidemiological risk environment where infection risk declines at a constant rate from $i_r = 0.6$ in age class 1 to $i_r = 0.2$ in age class 10. Reproduction begins in the third age class for all schedules. Strategy optima are determined as the immune specificity and sensitivity maximizing λ, the population growth rate. Dashed curve shows the shape of the specificity/sensitivity trade-off curve for γ = 4. Other parameter values are $\mu_b = 0.15$, $\mu_i = 0.1$, $\mu_d = 0.3$, and $\mu_{id} = 0.01$.

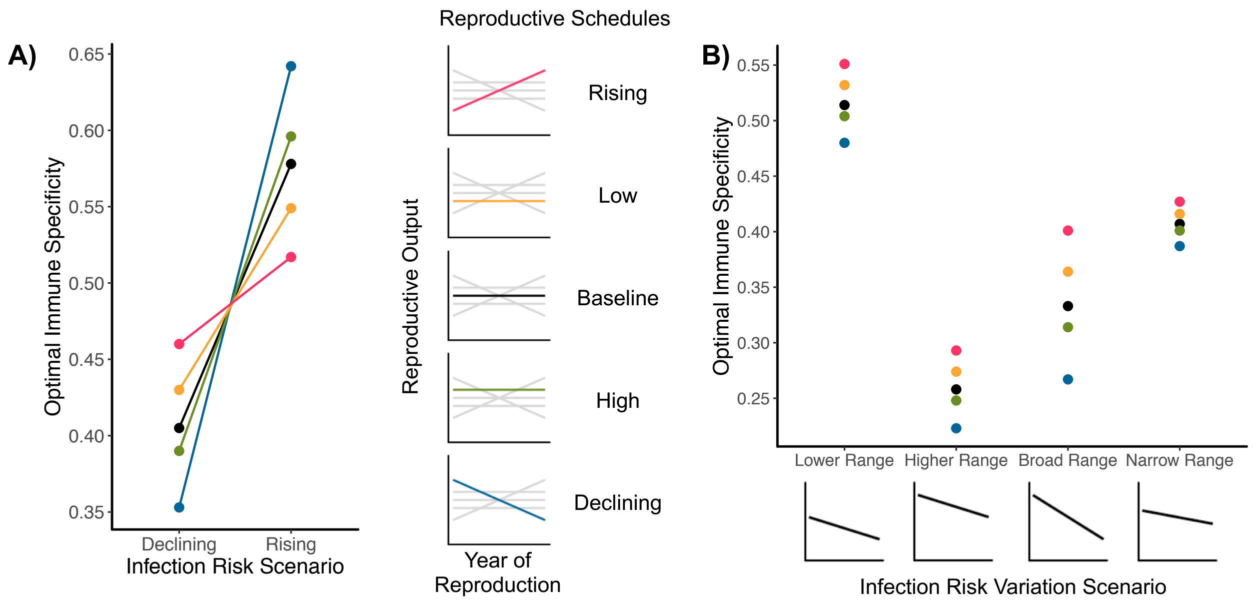


**Supplementary Figure 6: Epidemiological environment alters the effect of reproduction on immune strategy: infection risk $i_r$ with smoothed risk variation.** Reproduction begins in the third age class for all schedules, and $i_r$ changes a constant amount from age class to age class within each schedule. A) The change in optimal immune specificity associated with differences in epidemiological context (i.e. changes in $i_r$, on the x-axis) and reproduction (different points and lines). Parameter values are $\mu_b = 0.15$, $\mu_i = 0.1$, $\mu_d = 0.3$, $\mu_{id} = 0.01$, and $\gamma = 4$. In the declining scenario, $i_r$ declines with age from 0.6 to 0.2; in the rising scenario, $i_r$ increases from 0.2 to 0.6. B) The change in range of optimal specificities associated with different reproductive demographies associated with different magnitudes of variation in decline of infection risk $i_r$ with age. Parameter values are $\mu_b = 0.15$, $\mu_i = 0.1$, $\mu_d = 0.3$, $\mu_{di} = 0.01$, and $\gamma = 4$. In the lower range scenario, $i_r$ declines with age from 0.45 to 0.2; in the higher range, from 0.7 to 0.45; in the broad range, from 0.7 to 0.2; in the narrow range, from 0.525 to 0.375.

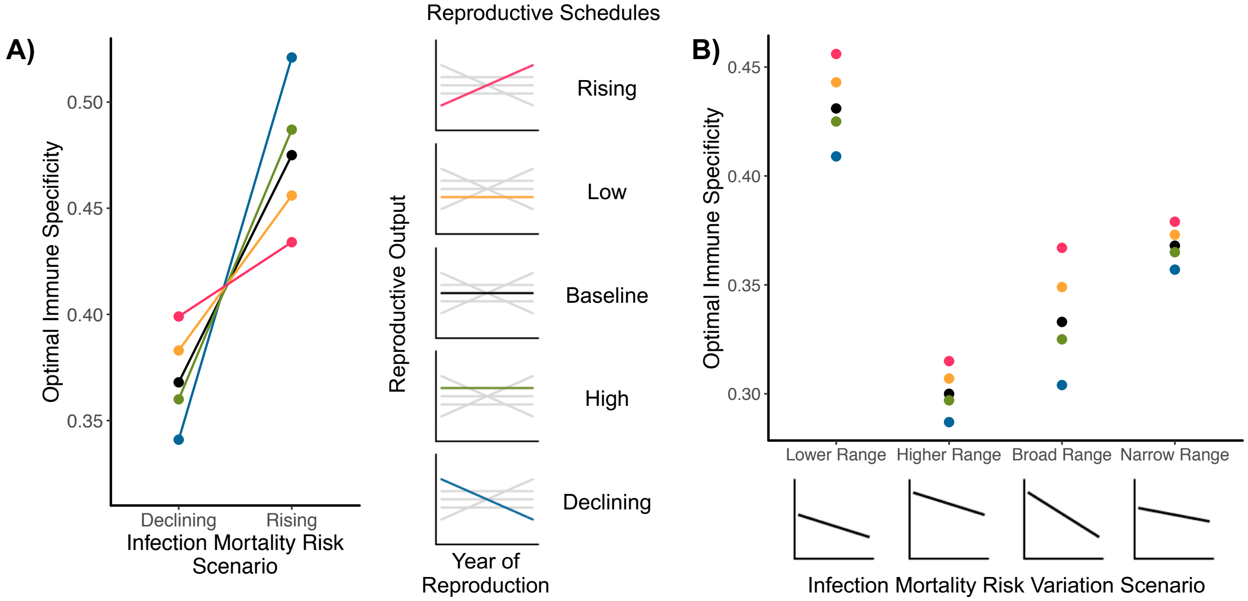


**Supplementary Figure 7: Epidemiological environment alters the effect of reproduction on immune strategy: infection mortality risk $\mu_d$ with smoothed risk variation.** Reproduction begins in the third age class for all schedules , and $\mu_d$ changes a constant amount from age class to age class within each schedule. A) The change in optimal immune specificity associated with differences in epidemiological context (i.e. changes in $\mu_d$, on the x-axis) and reproduction (different points and lines). Parameter values are $\mu_b = 0.15$, $\mu_i = 0.1$, $\mu_{id} = 0.01$, $i_r = 0.4$, and $\gamma = 4$. In the declining scenario, $\mu_d$ declines with age from 0.6 to 0.2; in the rising scenario, $\mu_d$ increases from 0.2 to 0.6. B) The change in range of optimal specificities associated with different reproductive demographies associated with different magnitudes of variation in decline of infection mortality risk $\mu_d$ with age. Parameter values are $\mu_b = 0.15$, $\mu_i = 0.1$, $\mu_{id} = 0.01$, $i_r = 0.4$, and $\gamma = 4$. In the lower range scenario, $\mu_d$ declines with age from 0.45 to 0.2; in the higher range, from 0.7 to 0.45; in the broad range, from 0.7 to 0.2; in the narrow range, from 0.525 to 0.375.

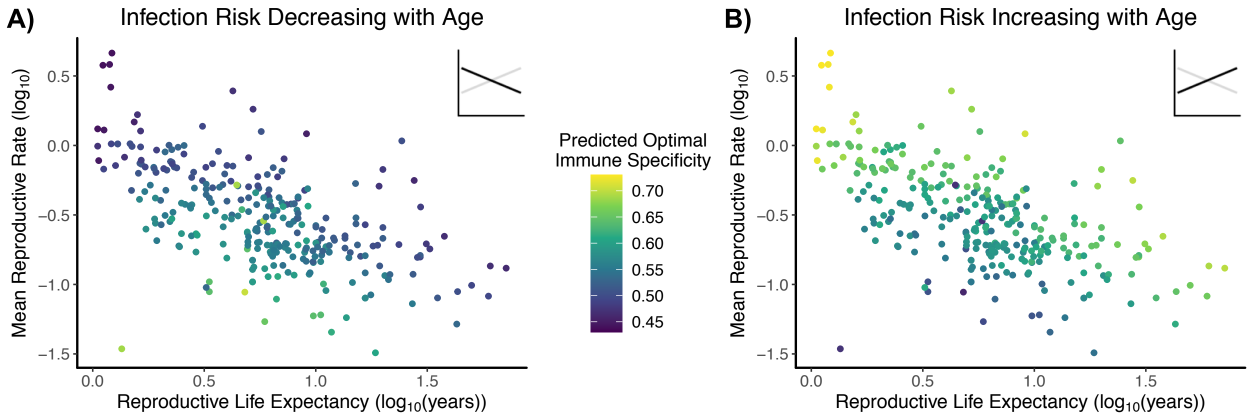


**Supplementary Figure 8: Interaction of demography and epidemiology: smoothed infection risk $i_r$ variation.** Infection risk is set such that infection risk changes at a constant rate relative to lifespan from a defined value of $i_r$ for the first age class to a defined value for the last age class. Our dataset comprises 298 population matrices representing 129 chordate species. For all scenarios, parameter values are $\mu_d$ = 0.3, $\mu_i$ = 0.1, $\mu_{id}$ = 0.01, $\rho$ = 0.75, and $\gamma$ = 4. A) Predicted optimal immune specificities when infection risk declines with age, with respect to population reproductive life expectancy and mean reproductive rate as calculated from original matrix. Infection risk $i_r$ prior to reproductive maturity is 0.45; for reproductive age classes, it is 0.2. B) Predicted optimal immune specificities when infection risk rises with age with respect to population reproductive life expectancy and mean reproductive rate as calculated from original matrix. Infection risk $i_r$ prior to reproductive maturity is 0.2; for reproductive age classes, it is 0.45.

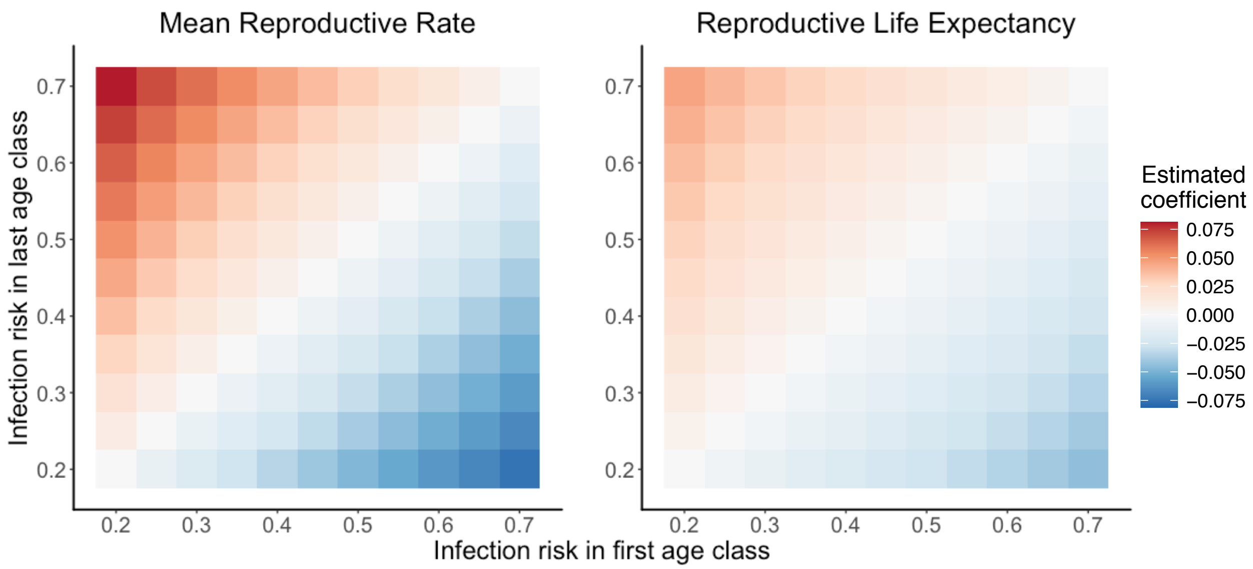


**Supplementary Figure 9: Epidemiological dependency of life history-specificity relationship: smoothed infection risk $i_r$ variation.** Tile plots showing, for a variety of scenarios of variation in $i_r$, coefficients of relationship between predicted optimal immune specificity and the designated life history trait, as estimated from Bayesian linear models. Each tile is a scenario in which predicted optimal immune specificities were generated for 298 population projection matrices representing 129 chordate species. A linear model was used to estimate relationship coefficients for each scenario. Coefficient value in plot represents mean value of the posterior probability distribution, except on the diagonal; on the diagonal, all coefficients are 0 (see text). Value of $i_r$ on x-axis is the risk for the first age class in the matrix, while the value on the y-axis is the risk for the last age class; rate of change between intermediate age classes is adjusted per matrix, based on dimension, so that risk changes at a constant rate from age class to age class within a matrix but the absolute magnitude of risk change is equivalent for all matrices. Life history trait values log-transformed and standardized as Z-scores for comparability of coefficients. For all estimated coefficients off the diagonal, 89% confidence intervals do not include 0. For all scenarios, parameter values are $\mu_d = 0.3$, $\mu_i = 0.1$, $\mu_{id} = 0.01$, $\rho = 0.75$, and $\gamma = 4$. Unlike Fig. 5, AFR is not shown because the model does not confidently predict a relationship with immune specificity for any $i_r$ scenario.

**Supplementary Table 1: Epidemiological Risk Scenarios: Set A.** The parameter values for each different analysis of the reproductive demography-immune strategy relationship as described in the Methods. When values are given in brackets, the first number is the parameter value in the first age class and the second number is the parameter value in the last age class of the matrix; for in-between age classes, parameter value either rises or falls in constant intervals from one age class to the next. We also considered similar scenarios in which $\mu_d$ varies on the same intervals as $i_r$ does here, while $i_r$ is held constant at 0.4.

| **Parameter** | **Scenario A1: Rising risk** | **Scenario A2: Declining risk** |
|---|---|---|
| $i_r$ | [0.6, 0.2] | [0.2, 0.6] |
| $\mu_b$ | 0.15 (identical for all scenarios) | |
| $\mu_i$ | 0.1 (identical for all scenarios) | |
| $\mu_d$ | 0.3 | 0.3 |
| $\mu_{id}$ | 0.01 (identical for all scenarios) | |
| $\gamma$ | 4 (identical for all scenarios) | |

**Supplementary Table 2: Epidemiological Risk Scenarios: Set B.** The parameter values for each scenario of the reproductive demography-immune strategy relationship as described in the Methods, used for an analysis of the influence of magnitude of change in risk with age on immune strategy. Reproductive maturity is in the third age class. We also considered scenarios where $\mu_d$ varied on the same intervals as $i_r$ does here, with $i_r$ then being held at 0.4 across all age classes.

| **Parameter** | **Scenario B1: Low range** | **Scenario B2: High range** | **Scenario B3: Broad range** | **Scenario B4: Narrow range** |
|---|---|---|---|---|
| **$i_r$ before reproductive maturity** | 0.45 | 0.7 | 0.7 | 0.525 |
| **$i_r$ after reproductive maturity** | 0.2 | 0.45 | 0.2 | 0.375 |
| $\mu_b$ | 0.15 (identical for all scenarios) | | | |
| $\mu_i$ | 0.1 (identical for all scenarios) | | | |
| $\mu_d$ | 0.3 (identical for all scenarios) | | | |
| $\mu_{id}$ | 0.01 (identical for all scenarios) | | | |
| $\gamma$ | 4 (identical for all scenarios) | | | |

**Supplementary Table 3: Effect of background mortality $\mu_b$ on optimal immune strategy.**

Comparison of results from analysis of effect of reproductive schedule on optimal immune strategy when $\mu_b$ does not change to equalize λ (original) and when $\mu_b$ does change to equalize λ (adjusted). $s_p^*$ is the optimal immune specificity that maximizes λ. Infection risk $i_r$ declines from 0.6 before reproductive maturity to 0.2 after. Other parameter values are $\mu_i = 0.1$, $\mu_d = 0.3$, $\mu_{id} = 0.01$, and $\gamma = 4$.

| **Reproductive Schedule** | **Original $\mu_b$** | **Original $\lambda_{max}$** | **Original $s_p^*$** | **Adjusted $\mu_b$** | **Adjusted $\lambda_{max}$** | **Adjusted $s_p^*$** |
|---|---|---|---|---|---|---|
| Rising | 0.150 | 1.089 | 0.590 | 0.105 | 1.133 | 0.591 |
| Low | 0.150 | 1.059 | 0.545 | 0.070 | 1.133 | 0.551 |
| Baseline | 0.150 | 1.133 | 0.517 | 0.150 | 1.133 | 0.517 |
| High | 0.150 | 1.193 | 0.497 | 0.216 | 1.133 | 0.493 |
| Declining | 0.150 | 1.199 | 0.439 | 0.224 | 1.133 | 0.436 |

**Supplementary Table 4: Effect of background mortality $\mu_b$ on optimal immune strategy: smoothed infection risk variation.** Comparison of results from analysis of reproductive schedule on optimal immune strategy when $\mu_b$ does not change to equalize λ (original) and when $\mu_b$ does change to equalize λ (adjusted). $s_p^*$ is the optimal immune specificity that maximizes λ. Infection risk $i_r$ declines smoothly from 0.6 in the first age class to 0.2 in the final age class. Other parameter values are $\mu_i$ = 0.1, $\mu_d$ = 0.3, $\mu_{id}$ = 0.01, and γ = 4.

| Reproductive Schedule | Original $\mu_b$ | Original $\lambda_{max}$ | Original $s_p^*$ | Adjusted $\mu_b$ | Adjusted $\lambda_{max}$ | Adjusted $s_p^*$ |
|---|---|---|---|---|---|---|
| Rising | 0.150 | 1.088 | 0.460 | 0.104 | 1.133 | 0.462 |
| Low | 0.150 | 1.059 | 0.430 | 0.070 | 1.133 | 0.436 |
| Baseline | 0.150 | 1.134 | 0.405 | 0.150 | 1.134 | 0.405 |
| High | 0.150 | 1.196 | 0.390 | 0.217 | 1.133 | 0.387 |
| Declining | 0.150 | 1.201 | 0.353 | 0.226 | 1.133 | 0.351 |

**Supplementary Table 5: Results from Bayesian linear model for demography and immune strategy – analysis using only mammal life histories.** Linear model looks at model-predicted optimal immune specificity as a function of three different life history summary statistics. Results are means and, in brackets, boundaries of 89% highest posterior density intervals. All variables calculated from original matrix in COMADRE database, log-transformed, and standardized as Z-scores. The only matrices included are those 151 qualifying matrices from 47 different mammal species. For stepped epidemiological scenario, when infection risk is rising, $i_r$ in pre-reproductive years is 0.2, and $i_r$ in reproductive years is 0.45. When infection risk declines in the stepped scenario, $i_r$ in pre-reproductive years is 0.45, and $i_r$ in reproductive years is 0.2. In smoothed declining scenario, $i_r$ declines from 0.45 to 0.2; in rising scenario, $i_r$ rises from 0.2 to 0.45. Other parameter values are $\mu_d$ = 0.3, $\mu_i$ = 0.1, $\mu_{id}$ = 0.01, ρ = 0.75, and γ = 4. *Values in italics* have confidence intervals overlapping with 0.

| Parameter | Declining stepped infection risk $i_r$ | Rising stepped infection risk $i_r$ | Declining smoothed infection risk $i_r$ | Rising smoothed infection risk $i_r$ |
|---|---|---|---|---|
| **Intercept** | 0.626 [0.624, 0.629] | 0.521 [0.518, 0.523] | 0.543 [0.540, 0.546] | 0.603 [0.599, 0.606] |
| **Age class of first reproduction** | -0.0407 [-0.0442, -0.0373] | 0.0356 [0.0326, 0.0384] | -0.00426 [-0.00830, $-3.74\times10^{-4}$] | *0.00311 [$-8.69\times10^{-4}$, 0.00699]* |
| **Mean reproductive rate** | -0.0309 [-0.0341, -0.0275] | 0.0266 [0.0239, 0.0293] | -0.0325 [-0.0363, -0.0285] | 0.0330 [0.0291, 0.0369] |
| **Reproductive life expectancy** | 0.00988 [0.00675, 0.0130] | -0.00894 [-0.0110, -0.00578] | -0.0313 [-0.0349, -0.0275] | 0.0342 [0.0306, 0.0379] |

| | | | | |
|---|---|---|---|---|
| **Standard deviation** | 0.0202 [0.0184, 0.0222] | 0.0176 [0.0160, 0.0194] | 0.0239 [0.0216, 0.0262] | 0.0247 [0.0224, 0.0271] |

**Supplementary Table 6: Results from Bayesian linear model for demography and immune strategy – analysis with generation time only.** Linear model looks at model-predicted optimal immune specificity as a function of the common life history summary statistic generation time, which is heavily confounded with other summary statistics. Results are means and, in brackets, boundaries of 89% highest posterior density intervals. All variables calculated from original matrix in COMADRE database, log-transformed, and standardized as Z-scores. Dataset includes 298 qualifying matrices from 129 chordate species. In smoothed declining scenario, $i_r$ declines from 0.45 to 0.2; in rising scenario, $i_r$ rises from 0.2 to 0.45. For stepped epidemiological scenario, when infection risk is rising, $i_r$ in pre-reproductive years is 0.2, and $i_r$ in reproductive years is 0.45. When infection risk declines in the stepped scenario, $i_r$ in pre-reproductive years is 0.45, and $i_r$ in reproductive years is 0.2. In smoothed declining scenario, $i_r$ declines from 0.45 to 0.2; in rising scenario, $i_r$ rises from 0.2 to 0.45. Other parameter values are $\mu_d = 0.3$, $\mu_i = 0.1$, $\mu_{id} = 0.01$, $\rho = 0.75$, and $\gamma = 4$.

| Parameter | Declining stepped infection risk $i_r$ | Rising stepped infection risk $i_r$ | Declining smoothed infection risk $i_r$ | Rising smoothed infection risk $i_r$ |
|---|---|---|---|---|
| **Intercept** | 0.603 [0.598, 0.608] | 0.544 [0.539, 0.548] | 0.538 [0.534, 0.542] | 0.608 [0.604, 0.611] |
| **Generation time** | 0.0268 [0.0221, 0.0315] | -0.0262 [-0.0308, -0.0216] | 0.0156 [0.0119, 0.0196] | -0.0168 [-0.0212, -0.0123] |
| **Standard deviation** | 0.0519 [0.0485, 0.0557] | 0.0503 [0.0471, 0.0540] | 0.0418 [0.0393, 0.0446] | 0.447 [0.0419, 0.0478] |

**Supplementary Table 7: Results from Bayesian linear model for demography and immune strategy – analysis with post-processing life history statistics.** Linear model looks at model-predicted optimal immune specificity as a function of three different life history summary statistics. Results are means and, in brackets, boundaries of 89% highest posterior density intervals. All variables calculated from output matrices after manipulation and optimization of $s_p$ (unlike results shown in Fig. 3 and Table 1), log-transformed, and standardized as Z-scores. Dataset includes 298 qualifying matrices from 129 chordate species. For stepped epidemiological scenario, when infection risk is declining, $i_r$ in pre-reproductive years is 0.2, and $i_r$ in reproductive years is 0.45. When infection risk declines in the stepped scenario, $i_r$ in pre-reproductive years is 0.45, and $i_r$ in reproductive years is 0.2. In smoothed declining scenario, $i_r$ declines from 0.45 to 0.2; in rising scenario, $i_r$ rises from 0.2 to 0.45. Other parameter values are $\mu_d$ = 0.3, $\mu_i$ = 0.1, $\mu_{id}$ = 0.01, ρ = 0.75, and γ = 4.

| **Parameter** | **Declining stepped infection risk $i_r$** | **Rising stepped infection risk $i_r$** | **Declining smoothed infection risk $i_r$** | **Rising smoothed infection risk $i_r$** |
|---|---|---|---|---|
| **Intercept** | 0.603 [0.600, 0.605] | 0.544 [0.541, 0.547] | 0.538 [0.536, 0.541] | 0.608 [0.605, 0.611] |
| **Age class of first reproduction** | -0.0379 [-0.0406, -0.0352] | 0.0359 [0.0333, 0.0386] | *$2.60x10^{-4}$ [-0.00304, 0.00351]* | *$-5.27x10^{-4}$ [-0.00387, 0.00300]* |
| **Mean reproductive rate** | -0.0420 [-0.0453, -0.0386] | 0.0407 [0.0374, 0.0439] | 0.0415 [-0.0452, -0.0376] | 0.0443 [0.0403, 0.0484] |
| **Reproductive life expectancy** | 0.0190 [0.0156, 0.0226] | -0.0193 [-0.0228, -0.0160] | -0.0223 [-0.0263, -0.0183] | 0.0236 [0.0194, 0.0277] |

| | | | | |
|---|---|---|---|---|
| **Standard deviation** | 0.0266<br>[0.0249, 0.0283] | 0.0261<br>[0.0243, 0.0279] | 0.0314<br>[0.0294, 0.0334] | 0.0334<br>[0.0313, 0.0356] |
