## Supplementary material for "Optimal immune specificity at the intersection of host life history and parasite epidemiology": R code used in analyses

# This script contains the R code for Downie et al. (2021).
# This first portion of the script is the functions needed to run the analyses.
# Some of the functions are original, while others are taken from either Metcalf and 
# Graham (2018) or Healy et al. (2019), and these latter will be marked.
# The second portion of the script contains the actual code for the analyses.

# If you have questions or concerns, please direct them to Alec Downie; I will be happy
# to help in any way that I can.

library(popbio)
library(popdemo)
library(MASS)
library(dplyr)
library(ggplot2)
library(rethinking)

### FUNCTIONS

##' Survival at age x as a function of specificity and other parameters
##' NOTE: This is directly from the code included with Metcalf and Graham (2018).
##'
##' @param mu.b background mortality hazard
##' @param mu.i immunopathology hazard
##' @param mu.d disease hazard
##' @param mu.di immunopathology and disease hazard
##' @param sp specificity
##' @param gamma parameter governing the efficiency of discrimination (high values indicate more discrimination)
##' @param ix probability of infection
##'
##' @return numeric
##'
sx <- function(mu.b,mu.i,mu.d,mu.di,sp=0.5, gamma=4, ix=0){
  exp(-(mu.b+
          mu.i*(1-sp)*(1-ix)+
          mu.d*exp(-gamma*(1-sp))*ix+
          mu.di*(1-exp(-gamma*(1-sp)))*ix))
}

##' Optimal specificity for a particular age
##' NOTE: This is directly from the code included with Metcalf and Graham (2018).
##'
##' @param mu.b background mortality hazard
##' @param mu.i immunopathology hazard
##' @param mu.d disease hazard
##' @param mu.di immunopathology and disease hazard
##' @param gamma parameter governing the efficiency of discrimination (high values indicate more discrimination)
##' @param ix probability of infection
##'
##' @return numeric
sx.max <- function(mu.b,mu.i,mu.d,mu.di,gamma=4, ix=0){
  1+(1/gamma)*(log(mu.i/(mu.d-mu.di))+log(1/gamma)+log((1-ix)/ix))
}

##' Make a Leslie matrix encompassing this probability of survival at every age
##' and with a chosen survival age trajectory
##' NOTE: This is directly from the code included with Metcalf and Graham (2018).
##'
##' @param mu.b background mortality hazard
##' @param mu.i immunopathology hazard
##' @param mu.d disease hazard
##' @param mu.di immunopathology and disease hazard
##' @param sp specificity
##' @param gamma parameter governing the efficiency of discrimination (high values indicate more discrimination)
##' @param ix a vector defining the probability of infection at each age - and note that its length defines the number of age classes
##' @param f.b a vector of fertility at every age, which should be the same length as ix
##'
##' @return list containing three vectors, survival only, fertility only, and the sum
##'
makeMat <- function(mu.b,mu.i,mu.d,mu.di,sp=0.5, gamma, ix, f.b) {
  
  nage <- length(ix)
  matU <- matF <- matrix(0,nage,nage)
  matF[1,] <- f.b
  
  psurv <- sx(mu.b=mu.b,mu.i=mu.i,mu.d=mu.d,mu.di=mu.di,sp=sp, gamma=gamma, ix=ix)
  matU[cbind(2:nage,1:(nage-1))] <- psurv[-length(psurv)]
  matU[nage,nage] <- psurv[length(psurv)]
  
  return(list(matU=matU,matF=matF,matM=matU+matF))
  
}

##' Expand a Leslie matrix to any size larger than its present size
##' 
##' @param matA a Leslie matrix of dimension n x n
##' @param dim a number denoting the dimensions of the desired new matrix
##' 
##' @return a matrix of the desired dimension expanded from the input
##' 
expandMat <- function(matA,dim){
  if (nrow(matA)!=ncol(matA)) stop("The input matrix must be square.")
  if (dim<=nrow(matA)) stop("You didn't request a matrix larger than the input one.")
  newMat <- matrix(data=rep(0,dim^2),nrow=dim,ncol=dim)
  newMat[1,] <- c(matA[1,1:(ncol(matA)-1)],rep(matA[1,ncol(matA)],dim-(ncol(matA)-1)))
  for (i in 2:nrow(newMat)){
    if(i<=nrow(matA))
      newMat[i,i-1]=matA[i,i-1]
    else newMat[i,i-1]=matA[nrow(matA),ncol(matA)]
  }
  newMat[dim,dim]=matA[nrow(matA),ncol(matA)]
  return(newMat)
}

##' A function to collapse matrices to a smaller dimension, according to the method
##' devised by Hooley (2000).  In brief, all rows/columns with indices greater than the
##' desired dimension are collapsed into a single row/column, using stable age structure
##' to weight survival probabilities and fertilities.
##' The collapsed rows and columns are those recommended by Salguero-Gomez and Plotkin
##' (2010): those corresponding to the latest age classes in the matrix.
##' See Hooley (2000) for a fuller explanation of the method.
##' 
##' @param matA A Leslie matrix of dimension n x n, to be collapsed to a smaller size
##' @param outputDim The desired dimension of the output matrix; must be an integer
##' 
##' @return a new matrix of dimension dim x dim, collapsed from the original matrix
##' 
Hooley <- function(matA,outputDim){
  # A* = PAQ, where A* is the output
  if (nrow(matA)!=ncol(matA)) stop("The input matrix must be square.")
  if (nrow(matA)<=outputDim) stop("You didn't request a matrix smaller than the input one.")
  origDim <- nrow(matA)
  matP <- matrix(0,nrow=outputDim,ncol=origDim)
  for (i in 1:nrow(matP)){
    if (i<nrow(matP)) matP[i,i]=1
    if (i==nrow(matP)) matP[i,i:ncol(matP)]=1
  }
  weights <- rep(0,origDim)
  stableAge <- stable.stage(matA)
  for (i in 1:origDim){
    if (i<outputDim) next()
    if (i>=outputDim) weights[i]=stableAge[i]/sum(stableAge[outputDim:length(stableAge)])
  }
  matQ <- matrix(0,nrow=origDim,ncol=outputDim)
  for (i in 1:ncol(matQ)){
    if (i<ncol(matQ)) matQ[i,i]=1
    if (i==ncol(matQ)) matQ[,i] <- weights
  }
  Astar <- matP %*% matA %*% matQ
  return(Astar)
}

##' A function that returns probability of reaching reproductive age, the age at first
##' at first reproduction (given as La), and the life expectancy of an individual reaching
##' reproductive maturity.
##' NOTE: This is taken directly from the code included in Healy et al. (2019)
##' 
##' @param matU a matrix giving the survival transition probabilities for the population
##' @param matF a matrix with the fecundities of each age class for the population
##' 
##' @return a data frame containing the life above life history summary stats
##' 
lifeTimeRepEvents <- function(matU, matF, startLife = 1){
  
  #Function to determine probability of reaching reproduction, age at maturity and reproductive lifespan (Code adapted from H. Caswell's matlab code):
  
  uDim = dim(matU)[1]
  surv = colSums(matU)
  repLifeStages = colSums(matF)
  repLifeStages[which(repLifeStages>0)] = 1
  
  if(missing(matF) | missing(matU)){stop('matU or matF missing')}
  if(sum(matF,na.rm=T)==0){stop('matF contains only 0 values')}
  
  #Probability of survival to first reprod event
  Uprime = matU
  Uprime[,which(repLifeStages==1)] = 0
  Mprime = matrix(0,2,uDim)
  for (p in 1:uDim[1]) {
    if (repLifeStages[p]==1) Mprime[2,p] = 1 else
      Mprime[1,p] = 1-surv[p]
  }
  
  Bprime = Mprime%*%(ginv(diag(uDim)-Uprime))
  pRep = Bprime[2,startLife]
  
  out = data.frame(pRep = pRep)
  #Age at first reproduction (La; Caswell 2001, p 124)
  D = diag(c(Bprime[2,]))
  Uprimecond = D%*%Uprime%*%ginv(D)
  expTimeReprod = colSums(ginv(diag(uDim)-Uprimecond))
  
  La = expTimeReprod[startLife] # Note that La is age class of first reproduction
  out$La = La
  
  #Mean life expectancy conditional on entering the life cycle in the first reproductive stage
  firstRepLifeStage = min(which(repLifeStages==1))
  
  N = solve(diag(uDim[1])-matU)
  
  meanRepLifeExpectancy = colSums(N)[firstRepLifeStage]
  
  out$meanRepLifeExpectancy = meanRepLifeExpectancy
  
  #Life expectancy from mean maturity
  remainingMatureLifeExpectancy = colSums(N)[startLife]-La
  
  out$remainingMatureLifeExpectancy = remainingMatureLifeExpectancy
  
  return(out)
  
}

##' A function that calculates the mean reproductive rate weighted for stable age
##' structure of the population.
##' NOTE: This is taken directly from the code included in Healy et al. (2019)
##' 
##' @param matA a matrix with both survival and fecundity for each age class
##' @param matF a matrix with the fecundities of each age class for the population
##' 
##' @return a number indicating the mean reproductive rate of the population.

meanRepo <- function(matA, matF){
  
  N <- dim(matF)[1]
  stable_stage <- eigen.analysis(matA)$stable.stage
  repo_sum <- vector()
  for(j in 1:(N)){
    repo_sum[j] <- sum(matF[,j])
  }
  mean_repo_rate <- as.vector(repo_sum %*% stable_stage)
  return(mean_repo_rate)
}


##' A function for collecting matrix summary statistics for comparison with optimal
##' immune strategy as derived above.  Used for determining matrix characters for
##' matrices from our optimal specificity ascertainment method.
##' 
##' @param matA A Leslie matrix
##' @param optSp The immune specificity used to generate matA
##' 
##' @return a list with a whole bunch of matrix summary statistics
##' 
matStats <- function(matA,optSp){
  gamma <- 4 # Can change if so desired!
  
  matA <- matA
  matF <- splitA(matA)[[2]]
  matU <- splitA(matA)[[1]]
  optSp <- optSp
  optSe <- 1-exp(-gamma*(1-optSp))
  dim <- ncol(matA)
  lambda <- lambda(matA)
  repValue <- reproductive.value(matA)
  expandRepValue <- reproductive.value(expandMat(matA,dim=(3*nrow(matA))))
  ageStruc <- stable.stage(matA)
  genTime <- generation.time(matA)
  someStats <- lifeTimeRepEvents(matU,matF)
  La <- someStats$La # La is age at first reproduction
  repLifeExpec <- someStats$meanRepLifeExpectancy
  meanRepRate <- meanRepo(matA,matF) # see function above
  meanRepRateNonStbl <- mean(matF[1,matF[1,]>0]) # code/concept from Healy et al. (2019),
  # describing the mean reproductive rate when not weighted for stable age structure.
  return(list(matA=matA,matF=matF,matU=matU,optSp=optSp,optSe=optSe,dim=dim,
              lambda=lambda,repValue=repValue,ageStruc=ageStruc,genTime=genTime,La=La,
              repLifeExpec=repLifeExpec,meanRepRate=meanRepRate,
              meanRepRateNonStbl=meanRepRateNonStbl,expandRepValue=expandRepValue))
}


##' A function that predicts optimal immune strategies for matrices by the method used
##' in this paper.  It takes an input list of matrices with IDs for the COMADRE database
##' and uses our described method to predict optimal immune sensitivity and specificity.
##' The output includes life history statistics for both pre and post-manipulation
##' versions of the matrix.
##' 
##' @param IDlist A vector containing the indices in COMADRE for each matrix of interest
##' @param irStart A value indicating i.r in the first age class.
##' @param irStop A value indicating i.r in the last age class.
##' @param schedType A string indicating how infection risk varies with age; the only
##' accepted values are "stepped" and "smoothed"
##' 
##' @return A data frame containing predicted immune strategies and a variety of life
##' history statistics for each matrix.
##' 

immune.predict <- function(IDlist,irStart,irStop,schedType){
  # This builds the output data frame.
  # "orig" variables are calculated from the original matrix referenced in the 
  # COMADRE database.  Regular parameters are calculated from the optimized matrix as
  # determined through the looping process.
  optSp <- c(0)
  optSe <- c(0)
  dim <- c(0)
  lambda <- c(0)
  genTime <- c(0)
  La <- c(0) # Age at first reproduction; this shorthand comes from Healy et al. (2019)
  meanRepRate <- c(0)
  repLifeExpec <- c(0)
  meanRepRateNonStbl <- c(0)
  origDim <- c(0)
  origLambda <- c(0)
  origGenTime <- c(0)
  origLa <- c(0)
  origMeanRepRate <- c(0)
  origRepLifeExpec <- c(0)
  origMeanRepRateNonStbl <- c(0)
  analysisResults <- data.frame(optSp,optSe,dim,lambda,genTime,La,
                                meanRepRate,repLifeExpec,meanRepRateNonStbl,
                                origDim,origLambda,origGenTime,origLa,origMeanRepRate,
                                origRepLifeExpec,origMeanRepRateNonStbl)
  
  
  # This next section does the actual process of ascertaining optimal strategies.
  for (k in 1:length(IDlist)){
    selectedMat <- comadre$mat[[IDlist[k]]]
    # The function below is taken from Healy et al. (2019)
    baseMatStats <- lifeTimeRepEvents(selectedMat$matU,selectedMat$matF)
    
    # This subsection does the matrix dimension manipulation
    scheduleLength <- baseMatStats$La+baseMatStats$meanRepLifeExpectancy
    origDim <- nrow(selectedMat$matA)
    
    if(ceiling(scheduleLength)==origDim) newMat <- selectedMat$matA
    if(ceiling(scheduleLength)>origDim){
      newMat <- expandMat(selectedMat$matA,ceiling(scheduleLength))
      # see the 'expandMat' function for more details
    }
    if(ceiling(scheduleLength)<origDim){
      newMat <- Hooley(selectedMat$matA,ceiling(scheduleLength))
      # see the 'Hooley' function for more details
    }
    
    # This subsection defines the infection schedules and other risk parameters.
    # These can be changed if you are interested in other scenarios.
    srvvl <- rep(0,ncol(newMat))
    for (i in 1:ncol(newMat)){
      if (i!=ncol(newMat)) srvvl[i]=newMat[i+1,i]
      if (i==ncol(newMat)) srvvl[i]=newMat[i,i]
    }
    rho <- 3/4
    mu.b <- rho*-log(srvvl)
    mu.d <- 0.3 # The mu.* values are from the Metcalf and Graham (2018) baseline.
    mu.i <- 0.1
    mu.di <- 0.01
    gamma <- 4
    
    if(schedType=="stepped"){
      ir <- rep(irStop,nrow(newMat))
      ir[1:(baseMatStats$La-1)] <- irStart
    }
    if(schedType=="smoothed"){
      ir <- seq(irStart,irStop,length.out=ncol(newMat))
    }
    if(schedType!="stepped"&schedType!="smoothed"){
      print("The parameter schedType must be either 'stepped' or 'smoothed'")
      return(NULL)
    }
    
    # This subsection identifies the optimal immune strategy (as defined above).
    sp <- seq(0,1,by=0.001)
    se <- 1-exp(-gamma*(1-sp))
    lambdas <- rep(0,length(sp))
    testMat <- matrix(data=rep(0,nrow(newMat)*ncol(newMat)),
                      nrow=nrow(newMat),ncol=ncol(newMat))
    testMat[1,] <- newMat[1,]
    for (i in 1:length(sp)){
      surv <- sx(mu.b,mu.i,mu.d,mu.di,sp=sp[i],gamma=4,ir)
      for (j in 1:ncol(newMat)){
        if (j<ncol(newMat)) {testMat[j+1,j]=surv[j]}
        if (j==ncol(newMat)) {testMat[j,j]=surv[j]}
      }
      lambdas[i]=lambda(testMat)
    }
    index <- which.max(lambdas)
    optSp <- sp[index]
    
    # This subsection reconstitutes the matrix with that optimal strategy and calculates
    # the life history summary statistics for said matrix.
    surv <- sx(mu.b,mu.i,mu.d,mu.di,sp=optSp,gamma=4,ir)
    optMat <- matrix(data=rep(0,nrow(newMat)*ncol(newMat)),
                     nrow=nrow(newMat),ncol=ncol(newMat))
    optMat[1,] <- newMat[1,]
    for (i in 1:ncol(newMat)){
      if (i<ncol(newMat)) {optMat[i+1,i]=surv[i]}
      if (i==ncol(newMat)) {optMat[i,i]=surv[i]}
    }
    
    origDetails <- matStats(selectedMat$matA,optSp=0.5) # optSp does not matter here.
    matDetails <- matStats(optMat,optSp)
    analysisResults[k,] <- c(matDetails$optSp,matDetails$optSe,matDetails$dim,
                             matDetails$lambda,
                             matDetails$genTime,matDetails$La,matDetails$meanRepRate,
                             matDetails$repLifeExpec,matDetails$meanRepRateNonStbl,
                             origDetails$dim,origDetails$lambda,origDetails$genTime,
                             origDetails$La,origDetails$meanRepRate,
                             origDetails$repLifeExpec,origDetails$meanRepRateNonStbl)
  }
  return(analysisResults)
}


### ANALYSES
# The code below creates Figure 2, and can be adapted to create Figure 3 as well as
# Figure S3-S6.

dim <- 10 # This sets the matrix size.  The risk parameters are arbitrary.
mu.b <- 0.15
mu.i <- 0.1
mu.d <- 0.3
mu.di <- 0.01
i.r <- rep(0.6,length.out=dim) # Infection risk declines (declining i.x)
gamma <- 4


# These vectors will form our data table
optSp <- rep(0,5)
optSe <- rep(0,5)
Reproduction <- rep(0,5)
optLambda <- rep(0,5)

# Here's where the reproductive schedule is defined
aFR <- 3 # age at first reproduction; the age class in which reproduction first occurs.
f.b <- rep(0,dim)
f.b[aFR:dim] <- rep(0.6,dim+1-aFR)
i.r[aFR:dim] <- rep(0.2,dim+1-aFR) # drop in i.r at reproductive maturity

spMax <- sx.max(mu.b,mu.i,mu.d,mu.di,gamma=4,ix=i.r) # Optimal specificity for each age

sp <- seq(0,1,by=0.001) # Set of specificities to be searched
se <- 1-exp(-gamma*(1-sp))
lambdas <- rep(0,length(sp))

for (i in 1:length(sp)){
  matB <- makeMat(mu.b,mu.i,mu.d,mu.di,sp=sp[i],gamma,i.r,f.b)
  lambdas[i] <- lambda(matB$M)
}
optSp[1] <- sp[which.max(lambdas)]
optSe[1] <- se[which.max(lambdas)]
Reproduction[1] <- "Baseline"
optLambda[1] <- max(lambdas)
# For more detailed demographic information
optMat <- makeMat(mu.b,mu.i,mu.d,mu.di,sp=optSp[1],gamma,i.r,f.b)
elstcMat <- elasticity(optMat$M)
for (i in 1:length(srvvlElstcty)){
  if (i<length(srvvlElstcty)){
    srvvlElstcty[i]=elstcMat[i+1,i]
  }
  if (i==length(srvvlElstcty)){
    srvvlElstcty[i]=elstcMat[i,i]
  }
}
frtltyElstcty <- elstcMat[1,]
elstctyInfo <- data.frame(ages,srvvlElstcty,frtltyElstcty,reproductive.value(optMat$M),
                          stable.stage(optMat$M))

# The next schedule is the one with raised (high) reproductive output
f.b[aFR:dim] <- rep(0.8,dim+1-aFR) # Change the reproductive schedule
lambdas <- rep(0,length(sp)) # Reset the fitness values
# mu.b <- 0.216 # These values allow recreation of Table S3.
for (i in 1:length(sp)){
  matB <- makeMat(mu.b,mu.i,mu.d,mu.di,sp=sp[i],gamma,i.r,f.b)
  lambdas[i] <- lambda(matB$M)
}
optSp[2] <- sp[which.max(lambdas)]
optSe[2] <- se[which.max(lambdas)]
Reproduction[2] <- "High"
optLambda[2] <- max(lambdas)
# For more detailed demographic information
optMat <- makeMat(mu.b,mu.i,mu.d,mu.di,sp=optSp[5],gamma,i.r,f.b)
elstcMat <- elasticity(optMat$M)
for (i in 1:length(srvvlElstcty)){
  if (i<length(srvvlElstcty)){
    srvvlElstcty[i]=elstcMat[i+1,i]
  }
  if (i==length(srvvlElstcty)){
    srvvlElstcty[i]=elstcMat[i,i]
  }
}
frtltyElstcty <- elstcMat[1,]
temp <- data.frame(ages,srvvlElstcty,frtltyElstcty,reproductive.value(optMat$M),
                   stable.stage(optMat$M))
elstctyInfo <- rbind(elstctyInfo,temp)

# The next schedule is the one with decreased (low) reproductive output
f.b[aFR:dim] <- rep(0.4,dim+1-aFR) # Change the reproductive schedule
lambdas <- rep(0,length(sp)) # Reset the fitness values
# mu.b <- 0.070
for (i in 1:length(sp)){
  matB <- makeMat(mu.b,mu.i,mu.d,mu.di,sp=sp[i],gamma,i.r,f.b)
  lambdas[i] <- lambda(matB$M)
}
optSp[3] <- sp[which.max(lambdas)]
optSe[3] <- se[which.max(lambdas)]
Reproduction[3] <- "Low"
optLambda[3] <- max(lambdas)
# For more detailed demographic information
optMat <- makeMat(mu.b,mu.i,mu.d,mu.di,sp=optSp[5],gamma,i.r,f.b)
elstcMat <- elasticity(optMat$M)
for (i in 1:length(srvvlElstcty)){
  if (i<length(srvvlElstcty)){
    srvvlElstcty[i]=elstcMat[i+1,i]
  }
  if (i==length(srvvlElstcty)){
    srvvlElstcty[i]=elstcMat[i,i]
  }
}
frtltyElstcty <- elstcMat[1,]
temp <- data.frame(ages,srvvlElstcty,frtltyElstcty,reproductive.value(optMat$M),
                   stable.stage(optMat$M))
elstctyInfo <- rbind(elstctyInfo,temp)

# The next schedule is the one with the rising reproductive output
f.b[aFR:dim] <- seq(0.1,1.1,length.out=dim+1-aFR) # Change the reproductive schedule
lambdas <- rep(0,length(sp)) # Reset the fitness values
# mu.b <- 0.105
for (i in 1:length(sp)){
  matB <- makeMat(mu.b,mu.i,mu.d,mu.di,sp=sp[i],gamma,i.r,f.b)
  lambdas[i] <- lambda(matB$M)
}
optSp[4] <- sp[which.max(lambdas)]
optSe[4] <- se[which.max(lambdas)]
Reproduction[4] <- "Rising"
optLambda[4] <- max(lambdas)
# For more detailed demographic information
optMat <- makeMat(mu.b,mu.i,mu.d,mu.di,sp=optSp[5],gamma,i.r,f.b)
elstcMat <- elasticity(optMat$M)
for (i in 1:length(srvvlElstcty)){
  if (i<length(srvvlElstcty)){
    srvvlElstcty[i]=elstcMat[i+1,i]
  }
  if (i==length(srvvlElstcty)){
    srvvlElstcty[i]=elstcMat[i,i]
  }
}
frtltyElstcty <- elstcMat[1,]
temp <- data.frame(ages,srvvlElstcty,frtltyElstcty,reproductive.value(optMat$M),
                   stable.stage(optMat$M))
elstctyInfo <- rbind(elstctyInfo,temp)

# The next schedule is the one with the declining reproductive output
f.b[aFR:dim] <- seq(1.1,0.1,length.out=dim+1-aFR) # Change the reproductive output
lambdas <- rep(0,length(sp)) # Reset the fitness values
# mu.b <- 0.224
for (i in 1:length(sp)){
  matB <- makeMat(mu.b,mu.i,mu.d,mu.di,sp=sp[i],gamma,i.r,f.b)
  lambdas[i] <- lambda(matB$M)
}
optSp[5] <- sp[which.max(lambdas)]
optSe[5] <- se[which.max(lambdas)]
Reproduction[5] <- "Declining"
optLambda[5] <- max(lambdas)
# For more detailed demographic information
optMat <- makeMat(mu.b,mu.i,mu.d,mu.di,sp=optSp[5],gamma,i.r,f.b)
elstcMat <- elasticity(optMat$M)
for (i in 1:length(srvvlElstcty)){
  if (i<length(srvvlElstcty)){
    srvvlElstcty[i]=elstcMat[i+1,i]
  }
  if (i==length(srvvlElstcty)){
    srvvlElstcty[i]=elstcMat[i,i]
  }
}
frtltyElstcty <- elstcMat[1,]
temp <- data.frame(ages,srvvlElstcty,frtltyElstcty,reproductive.value(optMat$M),
                   stable.stage(optMat$M))
elstctyInfo <- rbind(elstctyInfo,temp)

plotColors=c("black","#6B8E23","#FFA533","#FF3366","#006699")
# Those colors are black, green, orange, red, blue.
# They correspond to baseline, low, high, rising, declining.
results <- data.frame(optSp,optSe,Reproduction,optLambda,plotColors)

# Here's a plot of the results (Figure 2)
ggplot(tradeOff,aes(se,sp))+geom_line(color="grey10",size=0.75,linetype=2)+
  labs(x="Optimal Immune Sensitivity",y="Optimal Immune Specificity")+
  coord_cartesian(xlim=c(0.7,0.975),ylim=c(0.1,0.7))+
  scale_x_continuous(breaks=c(0.7,0.75,0.8,0.85,0.9,0.95))+
  scale_y_continuous(breaks=c(0.1,0.2,0.3,0.4,0.5,0.6,0.7))+
  theme_classic()+
  theme(axis.text=element_text(size=12),
        axis.line=element_line(size=0.75,lineend="round"),
        axis.title=element_text(size=14))+
  geom_hline(aes(yintercept=results$optSp[1]),color=plotColors[1],size=0.6)+
  geom_vline(aes(xintercept=results$optSe[1]),color=plotColors[1],size=0.6)+
  geom_hline(aes(yintercept=results$optSp[2]),color=plotColors[2],size=0.6)+
  geom_vline(aes(xintercept=results$optSe[2]),color=plotColors[2],size=0.6)+
  geom_hline(aes(yintercept=results$optSp[3]),color=plotColors[3],size=0.6)+
  geom_vline(aes(xintercept=results$optSe[3]),color=plotColors[3],size=0.6)+
  geom_hline(aes(yintercept=results$optSp[4]),color=plotColors[4],size=0.6)+
  geom_vline(aes(xintercept=results$optSe[4]),color=plotColors[4],size=0.6)+
  geom_hline(aes(yintercept=results$optSp[5]),color=plotColors[5],size=0.6)+
  geom_vline(aes(xintercept=results$optSe[5]),color=plotColors[5],size=0.6)

# To create Figure S3, you can simply plot each column of the elstctyInfo data frame
# against age.
elstctyInfo$Schedule <- rep(c("Baseline","High","Low","Rising","Declining"),each=10)
elstctyInfo$colors <- rep(plotColors,each=10)
elstctyInfo$Schedule <- factor(elstctyInfo$Schedule)
elstctyInfo <- rename(elstctyInfo,repValue=reproductive.value.optMat.M.,
                      stableAge=stable.stage.optMat.M.)

# To create Figure 3: loop the above analysis code through the six different infection
# risk scenarios below.  To speed the process up for Fig. 3 and Figs. S4-S7, you may wish 
# to skip the calculation of the detailed demographic information (reproductive values, 
# stable age structures, and elasticities).
# drop in i.r at reproductive maturity
i.r <- rep(0.6,length.out=dim) 
i.r[aFR:dim] <- rep(0.2,dim+1-aFR)
# rise in i.r at reproductive maturity
i.r <- rep(0.2,length.out=dim) 
i.r[aFR:dim] <- rep(0.6,dim+1-aFR)
# low range of i.r, dropping at reproductive maturity
i.r <- rep(0.45,length.out=dim) 
i.r[aFR:dim] <- rep(0.2,dim+1-aFR)
# high range of i.r, dropping at reproductive maturity
i.r <- rep(0.7,length.out=dim) 
i.r[aFR:dim] <- rep(0.45,dim+1-aFR)
# broad range of i.r, dropping at reproductive maturity
i.r <- rep(0.7,length.out=dim) 
i.r[aFR:dim] <- rep(0.2,dim+1-aFR)
# narrow range of i.r, dropping at reproductive maturity
i.r <- rep(0.525,length.out=dim) 
i.r[aFR:dim] <- rep(0.375,dim+1-aFR)

# To create Figure S4, you can use the same scenarios as above, but they should be for
# mu.d, rather than i.r.
# i.r should be a vector of the same length as the dimension of the matrix.  To repeat
# the analysis in the text, each value in the vector should be 0.4, but of course other
# values between 0 and 1 can be used.

# To create Figure S5, you can again use substantially the same code as above for Figure
# 2.  However, i.r needs to be defined as below for the smoothed decline.
i.r <- seq(0.6,0.2,length.out=dim) # Smoothed decline of i.r with age

# To create Figure S6, you can use the same procedure as for Figure 3, with the scenarios
# laid out below.  Each scenario is a smoothed version of the one used for Figure 3.
# decline in i.r with age
i.r <- seq(0.6,0.2,length.out=dim)
# rise in i.r with age
i.r <- seq(0.2,0.6,length.out=dim)
# low range decline in i.r with age
i.r <- seq(0.45,0.2,length.out=dim)
# high range decline in i.r with age
i.r <- seq(0.7,0.45,length.out=dim)
# broad range decline in i.r with age
i.r <- seq(0.7,0.2,length.out=dim)
# narrow range decline in i.r with age
i.r <- seq(0.525,0.375,length.out=dim)

# To create Figure S7, the procedure is much like that for Figure S4, where the scenarios
# are for changes in mu.d and i.r is held at 0.4 for all ages.


# The code below creates Figure 4 and Table 1
# FIRST, download the COMADRE database at https://compadre-db.org/Data/Comadre
# We used version 4.21.1.0.
# Then load the database from its directory into R; it will appear as "comadre"
# There is a helpful guide to its internal structure that can be found through the
# website linked above if you would like to know more about it.
species <- c()
goodMatNums <- c()

for (i in 1:length(comadre$metadata$SpeciesAccepted)){
  matA <- comadre$mat[[i]]$matA
  if (comadre$metadata$Phylum[i]!="Chordata"|
      is.na(comadre$metadata$Phylum[i])==TRUE) next # if you want to use all chordates
  # if (comadre$metadata$Class[i]!="Mammalia"|
  #     is.na(comadre$metadata$Class[i])==TRUE) next # if you want to use mammals only
  if (comadre$metadata$MatrixComposite[[i]]!="Mean"&
      comadre$metadata$MatrixComposite[[i]]!="Pooled") next 
  if (comadre$metadata$MatrixCriteriaAge[[i]]=="No"|
      is.na(comadre$metadata$MatrixCriteriaAge[[i]])==TRUE) next
  if (anyNA(matA)==TRUE) next # Turns out that some NAs crop up in COMADRE
  diagonal <- diag(matA)
  if (sum(diagonal[1:(length(diagonal)-1)])!=0) next
  if (isErgodic(matA)==FALSE) next
  if (isIrreducible(matA)==FALSE) next
  if (isPrimitive(matA)==FALSE) next
  goodMatNums=c(goodMatNums,i)
  species=c(species,comadre$metadata$SpeciesAccepted[i])
}
IDframe <- data.frame(species,goodMatNums)

# NOTE: depending on the number of matrices included in the input list, immune.predict()
# can take multiple minutes to run on a basic MacBook Pro.
analysisResultsIDstep <- immune.predict(IDframe$goodMatNums,0.45,0.2,"stepped")
analysisResultsIIstep <- immune.predict(IDframe$goodMatNums,0.2,0.45,"stepped")
analysisResultsIDsmooth <- immune.predict(IDframe$goodMatNums,0.45,0.2,"smoothed")
analysisResultsIIsmooth <- immune.predict(IDframe$goodMatNums,0.2,0.45,"smoothed")
# NOTE: if your two infection risk values are identical, then all matrices will have the
# same predicted optSp.  See Metcalf and Graham (2018) for the demonstration of this
# result (algebraic derivation).

# For the plots themselves
ggplot(analysisResultsIDstep,
       aes(origLa,log10(origMeanRepRate)))+
  geom_point(aes(color=optSp))+
  scale_color_viridis_c(limits=c(0.44,0.73),breaks=c(0.45,0.5,0.55,0.6,0.65,0.7))+
  theme_classic()+theme(axis.text=element_text(size=10),
                        axis.title=element_text(size=12),
                        plot.title=element_text(hjust=0.5,size=14),
                        axis.line=element_line(size=0.75,lineend="round"),
                        legend.title.align=0.5,
                        legend.text=element_text(angle=45),
                        legend.direction="horizontal")+
  labs(x="Age Class of First Reproduction",
       y=expression("Mean Reproductive Rate (log"[10]*")"),
       color="Model-Predicted Optimal\nImmune Specificity",
       title="Infection Risk Decreasing with Age")+guides(color="none")

ggplot(analysisResultsIIstep,
       aes(origLa,log10(origMeanRepRate)))+
  geom_point(aes(color=optSp))+
  scale_color_viridis_c(limits=c(0.44,0.73),breaks=c(0.45,0.5,0.55,0.6,0.65,0.7))+
  theme_classic()+theme(axis.text=element_text(size=10),
                        axis.title=element_text(size=12),
                        plot.title=element_text(hjust=0.5,size=14),
                        axis.line=element_line(size=0.75,lineend="round"),
                        legend.title.align=0.5,
                        legend.text.align=0.5)+
  labs(x="Age Class of First Reproduction",
       y=expression("Mean Reproductive Rate (log"[10]*")"),
       color="Model-Predicted Optimal\nImmune Specificity",
       title="Infection Risk Increasing with Age")+guides(color="none")

# Figure S8 can be adapted from the code above by switching the datasets used and the
# variable on the x-axis from AFR to reproductive life expectancy.

# For the analysis in the tables, first the variables of interest need to be standardized
analysisResultsIDstep$stdOrigLa <- scale(log(analysisResultsIDstep$origLa))
analysisResultsIDstep$stdOrigMeanRepRate <- scale(
  log(analysisResultsIDstep$origMeanRepRate))
analysisResultsIDstep$stdOrigRepLifeExpec <- scale(
  log(analysisResultsIDstep$origRepLifeExpec))

analysisResultsIIstep$stdOrigLa <- scale(log(analysisResultsIIstep$origLa))
analysisResultsIIstep$stdOrigMeanRepRate <- scale(
  log(analysisResultsIIstep$origMeanRepRate))
analysisResultsIIstep$stdOrigRepLifeExpec <- scale(
  log(analysisResultsIIstep$origRepLifeExpec))

analysisResultsIDsmooth$stdOrigLa <- scale(log(analysisResultsIDsmooth$origLa))
analysisResultsIDsmooth$stdOrigMeanRepRate <- scale(
  log(analysisResultsIDsmooth$origMeanRepRate))
analysisResultsIDsmooth$stdOrigRepLifeExpec <- scale(
  log(analysisResultsIDsmooth$origRepLifeExpec))

analysisResultsIIsmooth$stdOrigLa <- scale(log(analysisResultsIIsmooth$origLa))
analysisResultsIIsmooth$stdOrigMeanRepRate <- scale(
  log(analysisResultsIIsmooth$origMeanRepRate))
analysisResultsIIsmooth$stdOrigRepLifeExpec <- scale(
  log(analysisResultsIIsmooth$origRepLifeExpec))

predictSet <- dplyr::select(analysisResultsIDstep,optSp,stdOrigLa,
                            stdOrigMeanRepRate,stdOrigRepLifeExpec)
# You may wish to use more than 1 core for the model if your computer has multiple cores.
# Please note that, if your immune scenario features no variation in infection risk
# across age, then ulam() will return an error, because, as stated above, every optSp 
# value will be the same.
m.IDstep <- ulam(alist(
  optSp ~ dnorm(mu,sigma),
  mu <- a+b2*stdOrigLa+b3*stdOrigMeanRepRate+b4*stdOrigRepLifeExpec,
  a ~ dnorm(0.5,0.15),
  b2 ~ dnorm(0,0.1),
  b3 ~ dnorm(0,0.1),
  b4 ~ dnorm(0,0.1),
  sigma ~ dexp(1)
),data=predictSet,chains=4,cores=1,refresh=0)
precis(m.IDsimple,digits=4) # this will return a model summary

predictSet <- dplyr::select(analysisResultsIIstep,optSp,stdOrigLa,
                            stdOrigMeanRepRate,stdOrigRepLifeExpec)
m.IIstep <- ulam(alist(
  optSp ~ dnorm(mu,sigma),
  mu <- a+b2*stdOrigLa+b3*stdOrigMeanRepRate+b4*stdOrigRepLifeExpec,
  a ~ dnorm(0.5,0.15),
  b2 ~ dnorm(0,0.1),
  b3 ~ dnorm(0,0.1),
  b4 ~ dnorm(0,0.1),
  sigma ~ dexp(1)
),data=predictSet,chains=4,cores=1,refresh=0)
precis(m.IDsimple,digits=4)

predictSet <- dplyr::select(analysisResultsIDsmooth,optSp,stdOrigLa,
                            stdOrigMeanRepRate,stdOrigRepLifeExpec)
m.IDsmooth <- ulam(alist(
  optSp ~ dnorm(mu,sigma),
  mu <- a+b2*stdOrigLa+b3*stdOrigMeanRepRate+b4*stdOrigRepLifeExpec,
  a ~ dnorm(0.5,0.15),
  b2 ~ dnorm(0,0.1),
  b3 ~ dnorm(0,0.1),
  b4 ~ dnorm(0,0.1),
  sigma ~ dexp(1)
),data=predictSet,chains=4,cores=1,refresh=0)
precis(m.IDsimple,digits=4)

predictSet <- dplyr::select(analysisResultsIIsmooth,optSp,stdOrigLa,
                            stdOrigMeanRepRate,stdOrigRepLifeExpec)
m.IIsmooth <- ulam(alist(
  optSp ~ dnorm(mu,sigma),
  mu <- a+b2*stdOrigLa+b3*stdOrigMeanRepRate+b4*stdOrigRepLifeExpec,
  a ~ dnorm(0.5,0.15),
  b2 ~ dnorm(0,0.1),
  b3 ~ dnorm(0,0.1),
  b4 ~ dnorm(0,0.1),
  sigma ~ dexp(1)
),data=predictSet,chains=4,cores=1,refresh=0)
precis(m.IDsimple,digits=4)

# For Table S4, re-run the four prediction scenarios, but this time for a set of 
# matrices that only includes mammal taxa, not all chordates.  See the original
# filtering process, which contains the necessary code.  Then repeat the analyses as
# above, including with the standardization and shrinking of the data frame.  ulam()
# and the underlying Stan language can misbehave if the data frame contains columns
# that are not part of the model.

# For Table S5, use the model framework below after standardizing generation time and 
# building a small dataframe with just optSp and stdOrigGenTime, and run the model 
# across all four sets of predictions.
ulam(alist(
  optSp ~ dnorm(mu,sigma),
  mu <- a+b1*stdOrigGenTime,
  a ~ dnorm(0.5,0.15),
  b1 ~ dnorm(0,0.1),
  sigma ~ dexp(1)
),data=DATA,chains=4,cores=1,refresh=0)

# For Table S6, use the model framework below after standardizing the life history
# variable values from the post-processing matrices and building the corresponding 
# smaller data frame, and run the model across all four sets of predictions.
ulam(alist(
  optSp ~ dnorm(mu,sigma),
  mu <- a+b2*stdLa+b3*stdMeanRepRate+b4*stdRepLifeExpec,
  a ~ dnorm(0.5,0.15),
  b2 ~ dnorm(0,0.1),
  b3 ~ dnorm(0,0.1),
  b4 ~ dnorm(0,0.1),
  sigma ~ dexp(1)
),data=DATA,chains=4,cores=1,refresh=0)


# The code below generates Figure 5.
# Please be forewarned that this loop, if run on a basic computer, can take quite
# some time to run.  On the computer with which we did the analysis, it took ~4 hours.
# If you are interested in playing with the conditions, you may want to reduce the 
# number of infection scenarios you loop through by altering i.rFirst and i.rLast.
i.rFirst <- rep(seq(0.2,0.7,by=0.05),each=11)
i.rLast <- rep(seq(0.2,0.7,by=0.05),times=11)

# This builds all the columns that will go into our output data frame.
estIntercept <- rep(0,length(i.rFirst))
sdIntercept <- rep(0,length(i.rFirst))
HPDIlowIntercept <- rep(0,length(i.rFirst))
HPDIhighIntercept <- rep(0,length(i.rFirst))
HPDI95lowIntercept <- rep(0,length(i.rFirst))
HPDI95highIntercept <- rep(0,length(i.rFirst))
estAFR <- rep(0,length(i.rFirst))
sdAFR <- rep(0,length(i.rFirst))
HPDIlowAFR <- rep(0,length(i.rFirst))
HPDIhighAFR <- rep(0,length(i.rFirst))
HPDI95lowAFR <- rep(0,length(i.rFirst))
HPDI95highAFR <- rep(0,length(i.rFirst))
estMRR <- rep(0,length(i.rFirst))
sdMRR <- rep(0,length(i.rFirst))
HPDIlowMRR <- rep(0,length(i.rFirst))
HPDIhighMRR <- rep(0,length(i.rFirst))
HPDI95lowMRR <- rep(0,length(i.rFirst))
HPDI95highMRR <- rep(0,length(i.rFirst))
estRLE <- rep(0,length(i.rFirst))
sdRLE <- rep(0,length(i.rFirst))
HPDIlowRLE <- rep(0,length(i.rFirst))
HPDIhighRLE <- rep(0,length(i.rFirst))
HPDI95lowRLE <- rep(0,length(i.rFirst))
HPDI95highRLE <- rep(0,length(i.rFirst))
estSigma <- rep(0,length(i.rFirst))
sdSigma <- rep(0,length(i.rFirst))
HPDIlowSigma <- rep(0,length(i.rFirst))
HPDIhighSigma <- rep(0,length(i.rFirst))
HPDI95lowSigma <- rep(0,length(i.rFirst))
HPDI95highSigma <- rep(0,length(i.rFirst))

# This specifies the model for use in the loop.
m <- alist(
  optSp ~ dnorm(mu,sigma),
  mu <- a+b2*stdOrigLa+b3*stdOrigMeanRepRate+b4*stdOrigRepLifeExpec,
  a~dnorm(0.5,0.15),
  b2~dnorm(0,0.1),
  b3~dnorm(0,0.1),
  b4~dnorm(0,0.1),
  sigma~dexp(1)
)

for (i in (1:length(i.rFirst))){
  predictSet <- immune.predict(IDframe$goodMatNums,i.rFirst[i],i.rLast[i],"stepped")
  
  # Standardization of variables is needed.
  predictSet$stdOrigLa <- scale(log(predictSet$origLa))
  predictSet$stdOrigMeanRepRate <- scale(log(predictSet$origMeanRepRate))
  predictSet$stdOrigRepLifeExpec <- scale(log(predictSet$origRepLifeExpec))
  
  # The data frame should be pruned to only contain variables of interest.
  predictSet <- dplyr::select(predictSet,optSp,stdOrigLa,
                              stdOrigMeanRepRate,stdOrigRepLifeExpec)
  
  # The modeling function produces an error when infection risk does not change, 
  # because the optimal immune defense is constant.  Therefore these must be skipped.
  if (length(unique(predictSet$optSp))==1){
    next()
  } else {
    outputModel <- ulam(m,data=predictSet,chains=4,cores=1,refresh=0,iter=2000)
    modelPrecis89 <- precis(outputModel,digits=4,prob=0.89)
    modelPrecis95 <- precis(outputModel,digits=4,prob=0.95)
    # You may find that you are comfortable with just one highest posterior density
    # interval, rather than the two here.
    
    # Now we store all the outputs (clumsily).
    estIntercept[i] <- modelPrecis89[1,1]
    sdIntercept[i] <- modelPrecis89[1,2]
    HPDIlowIntercept[i] <- modelPrecis89[1,3]
    HPDIhighIntercept[i] <- modelPrecis89[1,4]
    HPDI95lowIntercept[i] <- modelPrecis95[1,3]
    HPDI95highIntercept[i] <- modelPrecis95[1,4]
    estAFR[i] <- modelPrecis89[2,1]
    sdAFR[i] <- modelPrecis89[2,2]
    HPDIlowAFR[i] <- modelPrecis89[2,3]
    HPDIhighAFR[i] <- modelPrecis89[2,4]
    HPDI95lowAFR[i] <- modelPrecis95[2,3]
    HPDI95highAFR[i] <- modelPrecis95[2,4]
    estMRR[i] <- modelPrecis89[3,1]
    sdMRR[i] <- modelPrecis89[3,2]
    HPDIlowMRR[i] <- modelPrecis89[3,3]
    HPDIhighMRR[i] <- modelPrecis89[3,4]
    HPDI95lowMRR[i] <- modelPrecis95[3,3]
    HPDI95highMRR[i] <- modelPrecis95[3,4]
    estRLE[i] <- modelPrecis89[4,1]
    sdRLE[i] <- modelPrecis89[4,2]
    HPDIlowRLE[i] <- modelPrecis89[4,3]
    HPDIhighRLE[i] <- modelPrecis89[4,4]
    HPDI95lowRLE[i] <- modelPrecis95[4,3]
    HPDI95highRLE[i] <- modelPrecis95[4,4]
    estSigma[i] <- modelPrecis89[5,1]
    sdSigma[i] <- modelPrecis89[5,2]
    HPDIlowSigma[i] <- modelPrecis89[5,3]
    HPDIhighSigma[i] <- modelPrecis89[5,4]
    HPDI95lowSigma[i] <- modelPrecis95[5,3]
    HPDI95highSigma[i] <- modelPrecis95[5,4]
    
    # A little memory clean-up.
    rm(predictSet)
    rm(outputModel)
  }
}

coefficientTable <- data.frame(i.rFirst,i.rLast,estIntercept,sdIntercept,
                               HPDIlowIntercept,HPDIhighIntercept,
                               HPDI95lowIntercept,HPDI95highIntercept,
                               estAFR,sdAFR,HPDIlowAFR,HPDIhighAFR,
                               HPDI95lowAFR,HPDI95highAFR,
                               estMRR,sdMRR,HPDIlowMRR,HPDIhighMRR,
                               HPDI95lowMRR,HPDI95highMRR,
                               estRLE,sdRLE,HPDIlowRLE,HPDIhighRLE,
                               HPDI95lowRLE,HPDI95highRLE,
                               estSigma,sdSigma,HPDIlowSigma,HPDIhighSigma,
                               HPDI95lowSigma,HPDI95highSigma)

# For the plots in the figure themselves.  Note that the range for the fill geom is
# set manually here.
ggplot(coefficientTable,aes(i.rFirst,i.rLast))+
  geom_raster(aes(fill=estMRR),interpolate=FALSE)+
  scale_fill_distiller(palette="RdBu",limits=c(-0.077,0.077),
                       breaks=c(-0.075,-0.05,-0.025,0.00,0.025,0.05,0.075))+
  scale_x_continuous(breaks=c(0.2,0.3,0.4,0.5,0.6,0.7))+
  scale_y_continuous(breaks=c(0.2,0.3,0.4,0.5,0.6,0.7))+
  theme_classic()+
  theme(axis.text=element_text(size=12),
        axis.line=element_line(size=0.75,lineend="round"),
        axis.title=element_text(size=14),
        plot.title=element_text(size=16,hjust=0.5),
        legend.title=element_text(hjust=0.5))+
  labs(x="Infection risk for pre-reproductive years",
       y="Infection risk for reproductive years",
       title="Mean Reproductive Rate",
       fill="Estimated\ncoefficient")

ggplot(coefficientTable,aes(i.rFirst,i.rLast))+
  geom_raster(aes(fill=estAFR),interpolate=FALSE)+
  scale_fill_distiller(palette="RdBu",limits=c(-0.077,0.077),
                       breaks=c(-0.075,-0.05,-0.025,0.00,0.025,0.05,0.075))+
  scale_x_continuous(breaks=c(0.2,0.3,0.4,0.5,0.6,0.7))+
  scale_y_continuous(breaks=c(0.2,0.3,0.4,0.5,0.6,0.7))+
  theme_classic()+
  theme(axis.text=element_text(size=12),
        axis.line=element_line(size=0.75,lineend="round"),
        axis.title=element_text(size=14),
        plot.title=element_text(size=16,hjust=0.5),
        legend.title=element_text(hjust=0.5))+
  labs(x="Infection risk for pre-reproductive years",
       y="Infection risk for reproductive years",
       title="Age Class of First Reproduction",
       fill="Estimated\ncoefficient")

ggplot(coefficientTable,aes(i.rFirst,i.rLast))+
  geom_raster(aes(fill=estRLE),interpolate=FALSE)+
  scale_fill_distiller(palette="RdBu",limits=c(-0.077,0.077),
                       breaks=c(-0.075,-0.05,-0.025,0.00,0.025,0.05,0.075))+
  scale_x_continuous(breaks=c(0.2,0.3,0.4,0.5,0.6,0.7))+
  scale_y_continuous(breaks=c(0.2,0.3,0.4,0.5,0.6,0.7))+
  theme_classic()+
  theme(axis.text=element_text(size=12),
        axis.line=element_line(size=0.75,lineend="round"),
        axis.title=element_text(size=14),
        plot.title=element_text(size=16,hjust=0.5),
        legend.title=element_text(hjust=0.5))+
  labs(x="Infection risk for pre-reproductive years",
       y="Infection risk for reproductive years",
       title="Reproductive Life Expectancy",
       fill="Estimated\ncoefficient")

# To make Figure S9, everything can stay the same, except immune.predict() should use
# the "smoothed" specification rather than "stepped."  You will want to alter the range
# for the coefficients in the plots, otherwise the tile coloring won't quite be right.
